# Mutations in the *Staphylococcus aureus* Global Regulator CodY Confer Tolerance to an Interspecies Redox-Active Antimicrobial

**DOI:** 10.1101/2024.07.02.601769

**Authors:** Anthony M. Martini, Sara A. Alexander, Anupama Khare

**Affiliations:** Laboratory of Molecular Biology, Center for Cancer Research, National Cancer Institute, National Institutes of Health, Bethesda, MD, USA

**Keywords:** *Staphylococcus aureus*, pyocyanin, antimicrobial tolerance, reactive oxygen species

## Abstract

Bacteria often exist in multispecies communities where interactions among different species can modify individual fitness and behavior. Although many competitive interactions have been characterized, molecular adaptations that can counter this antagonism and preserve or increase fitness remain underexplored. Here, we characterize the adaptation of *Staphylococcus aureus* to pyocyanin, a redox-active interspecies antimicrobial produced by *Pseudomonas aeruginosa*, a co-infecting pathogen frequently isolated from wound and chronic lung infections with *S. aureus*. Using experimental evolution, we identified mutations in a conserved global transcriptional regulator, CodY, that confer tolerance to pyocyanin and thereby enhance survival of *S. aureus*. The transcriptional response of a pyocyanin tolerant CodY mutant to pyocyanin indicated a two-pronged defensive response compared to the wild type. Firstly, the CodY mutant strongly suppressed metabolism, by downregulating pathways associated with core metabolism, especially translation-associated genes, upon exposure to pyocyanin. Metabolic suppression via ATP depletion was sufficient to provide comparable protection against pyocyanin to the wild-type strain. Secondly, while both the wild-type and CodY mutant strains upregulated oxidative stress response pathways, the CodY mutant overexpressed multiple stress response genes compared to the wild type. We determined that catalase overexpression was critical to pyocyanin tolerance as its absence eliminated tolerance in the CodY mutant and overexpression of catalase was sufficient to impart tolerance to the wild-type strain. Together, these results suggest that both transcriptional responses likely contribute to pyocyanin tolerance in the CodY mutant. Our data thus provide new mechanistic insight into adaptation toward interbacterial antagonism via altered regulation that facilitates multifaceted protective cellular responses.

## INTRODUCTION

Microorganisms commonly live in the presence of other microbial species, whether in diverse environmental niches or in association with a host (1–3). These polymicrobial communities can be structurally and functionally dynamic in part through the balance of cooperative and competitive interactions among members (4, 5). Through these molecular interactions, microbial species can impact the fitness, behaviors, and adaptation of other constituent members of the community (6–8). Notably, antimicrobial effects of several secreted compounds have been shown to mediate interbacterial antagonism *in vitro*, enhancing the relative fitness of the producing species (9, 10). How community members may adapt to these antagonistic interactions is, however, less well-characterized.

The potential role of microbial interactions in human disease is being increasingly appreciated (11). In particular, the prevalence of *Staphylococcus aureus* and *Pseudomonas aeruginosa* co-infection in wounds (12) and in the airways of people with cystic fibrosis (CF) (13) has prompted extensive work characterizing the molecular interactions between these two pathogens (14–16). In CF, simultaneous culture of *S. aureus* and *P. aeruginosa* is associated with more deleterious clinical characteristics compared to mono-infection with either pathogen in some cohorts (17–19) but not others (20–22). Interestingly, although *P. aeruginosa* rapidly eradicates *S. aureus* under typical *in vitro* conditions (15, 23), co-colonization with both pathogens *in vivo* can persist for years (24). This suggests that one or both species likely exhibit altered physiology and/or spatial partitioning *in vivo* and adaptations may further contribute to their co-existence.

*P. aeruginosa* virulence is largely attributable to the many toxins it produces (25). Among these toxins is the redox-active secondary metabolite, pyocyanin (PYO) (26, 27). PYO has been detected in secretions produced during ear infection (28) and the sputum and large airways of people with CF (29) where it can contribute to cellular toxicity (26, 30). In addition to virulence, PYO is known to have antimicrobial properties against several other microbial species via the production of reactive oxygen species or inhibition of the electron transport chain (ETC) (31–33).

Bacteria can adapt to the presence of antimicrobials by evolving resistance, where they can grow in higher concentrations of the antimicrobial, or tolerance, where they survive in higher concentrations of the antimicrobial (34, 35). Previous studies have identified adaptations leading to PYO resistance in *Escherichia coli*, likely by reducing intracellular PYO concentrations and altering metabolism (36), and in *Agrobacterium tumefaciens* by altering electron transport chain function and increasing the oxidative stress response, although other mechanisms likely also play a role (37). In *S. aureus*, it has been shown that PYO resistance can be conferred by mutations in respiratory chain components and via putative quinone resistance responses (38–40), but additional mechanisms, especially of PYO tolerance, remain unknown.

In this study we investigate the ability of *S. aureus* to adapt to the bactericidal effects of PYO using experimental evolution, identifying novel mechanisms of PYO tolerance. In our evolved populations and isolates, we observe ubiquitous mutations in CodY, a conserved transcriptional regulator of virulence and metabolism in gram-positive bacteria (41). The breadth of mutations observed in *codY* are likely to reduce CodY activity and we show that CodY loss-of-function confers enhanced survival during treatment with PYO. Transcriptional analysis indicates a strong response to reactive oxygen stress during PYO treatment in both the wild-type and a CodY mutant. However, we observe that loss of CodY activity both suppresses translation-associated gene expression and produces a stronger oxidative stress response compared to WT. Finally, we demonstrate that recapitulating these phenotypes individually via metabolic suppression through ATP depletion or the overexpression of hydrogen peroxide-detoxifying catalase, protect WT cells from PYO-mediated cell death, indicating that both these mechanisms likely underlie the enhanced PYO survival of the *codY\** mutant. Thus, mutations in a global regulator can fine-tune the regulatory landscape to enable a multidimensional adaptive response to interspecies toxins.

## RESULTS

### Experimental evolution selects for pyocyanin tolerance in *S. aureus*

We first determined the bactericidal effect of PYO on *S. aureus* strain JE2 by quantifying survival of exponential phase cells upon treatment with a range of PYO concentrations. We observed a concentration-dependent effect of PYO on *S. aureus* cell density, including moderate growth reduction at 12.5 and 25 µM, growth inhibition at 50 and 100 µM, and killing at 200 and 400 µM (**Fig. 1A**).

**Figure 1.**
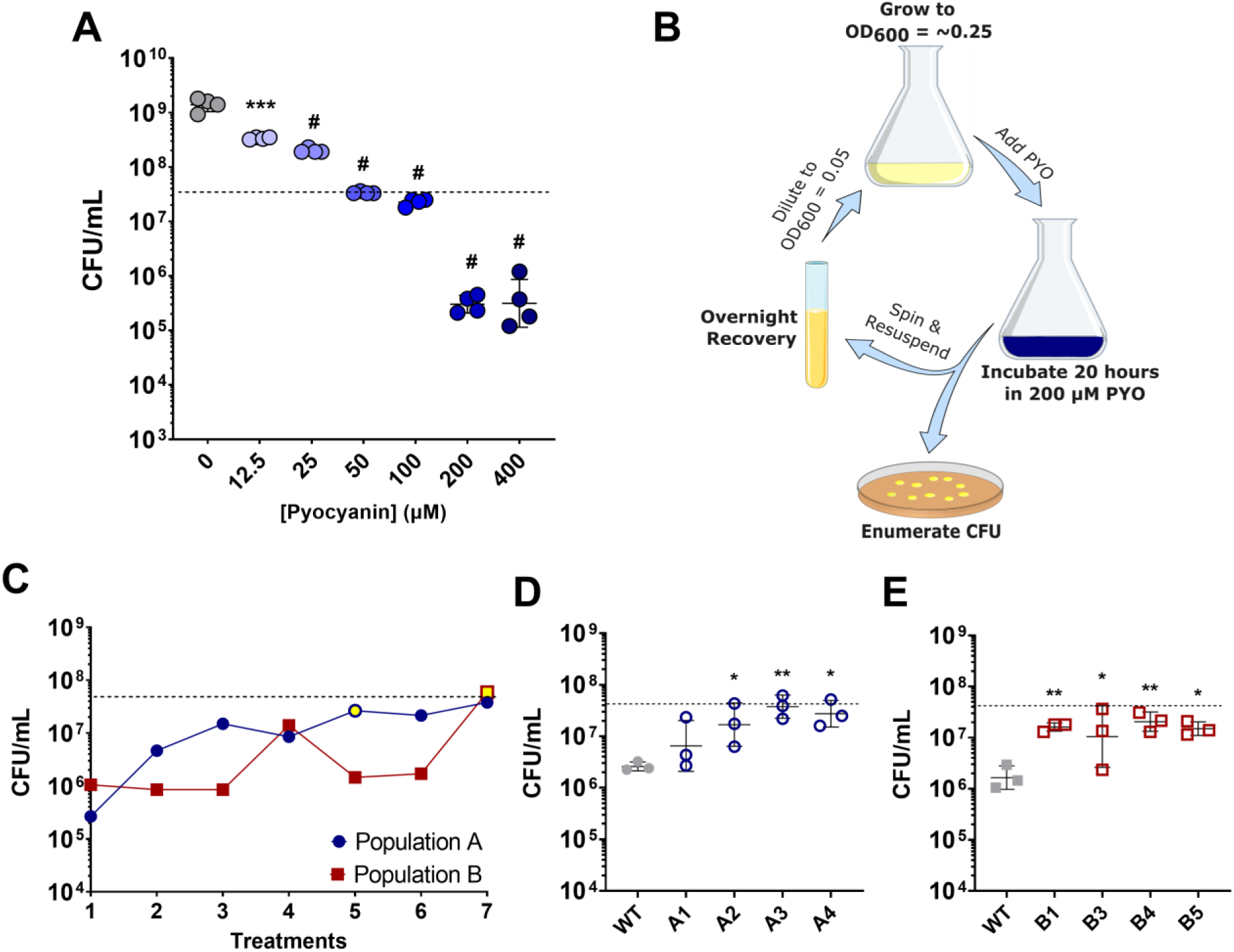
Experimental evolution of *S. aureus* selects for enhanced survival in PYO. (**A**) Early exponential phase *S. aureus* treated with the indicated concentration of PYO. Values indicate *S. aureus* viable cell counts after 20 hours of PYO exposure. Data shown are the geometric mean ± geometric standard deviation of four biological replicates. (**B**) Schematic of experimental evolution to select for PYO-tolerant *S. aureus*. Overnight cultures were grown to early exponential-phase prior to addition of 200 µM PYO. After incubation for 20 hours, viable cell counts were enumerated, and the remaining culture was grown overnight in the absence of PYO. This process was repeated six times for two independent populations. (**C**) Viable cell counts of two independent populations were experimentally evolved, and the viable cell counts monitored after each treatment with 200 µM PYO. Yellow fill indicates the passage at which isolates were selected. (**D, E**) Selected isolates from the indicated populations were assayed as in (**A**) for tolerance to 200 µM PYO. Data shown are the geometric mean ± geometric standard deviation of three biological replicates. (**A, C, D, E**) The dashed line indicates the mean initial cell density (CFU/mL) for all strains at the time of addition of PYO or DMSO as a control (0 µM PYO). Significance is indicated for comparison to the DMSO control (0 µM PYO) (**A**), or WT (**D, E**) as determined by a one-way ANOVA using Dunnett’s correction for multiple comparisons (\**P* < 0.05, \*\**P* < 0.01, \*\*\**P* < 0.001, ^#^*P* < 0.0001).

To identify potential adaptations that increase survival of *S. aureus* upon PYO exposure, we decided to use experimental evolution of cells upon repeated exposures to 200 µM PYO – the lowest bactericidal concentration identified. We evolved two independent populations by treating early exponential phase cells with PYO for 20 hours, recovering the surviving cells overnight in media, and repeating this process over several iterations (**Fig. 1B**). Because we observed loss of cell viability with this concentration of PYO and recovered surviving cells, our expectation was that we would identify mutants exhibiting increased survival during treatment with PYO. Indeed, as the number of treatments increased, we observed enhanced survival of both independent populations when treated with PYO (**Fig. 1C**). Individual isolates from the evolved populations also exhibited higher survival following PYO exposure, indicating that these evolved strains acquired increased tolerance to *P. aeruginosa*-derived PYO (**Fig. 1D, 1E** and **Supp. Fig. 1**). The selection of strains exhibiting reduced cell death, rather than a continuation of growth, during treatment with PYO suggested that we were selecting for PYO tolerance, rather than resistance.

### Loss-of-function mutations in the CodY global regulator confer tolerance to pyocyanin

Next, we sought to identify common mutations that characterize PYO-tolerant isolates from each evolved population, and thus sequenced and analyzed genomes from 18 terminal isolates (11 isolates from population A and 7 isolates from population B). While we observed diverse mutations among different isolates (**Supp. Data File 01**), each of the 18 isolates had at least one mutation associated with the *codY* gene (**Supp. Table 1**), which encodes a well-characterized pleiotropic transcriptional regulator conserved across gram-positive bacteria (41). In *S. aureus*, CodY regulates the expression of virulence and metabolic genes in response to nutritional cues (42, 43); however, a role in modulating tolerance to interspecies antimicrobials has not, to our knowledge, been described. Coding sequence mutations observed in our evolved isolates were present in both the substrate sensing and DNA-binding domains, and we also observed mutations in the promoter region and the start codon (**Fig. 2A** and **Supp. Table 1**). Isolation of the R61K mutation, which has previously been described to substantially reduce CodY activity (42), and an ablated start-codon, as well as the diversity of mutations across both functional domains suggested that the mutations we observed in our evolved isolates likely resulted in loss of CodY function.

**Figure 2.**
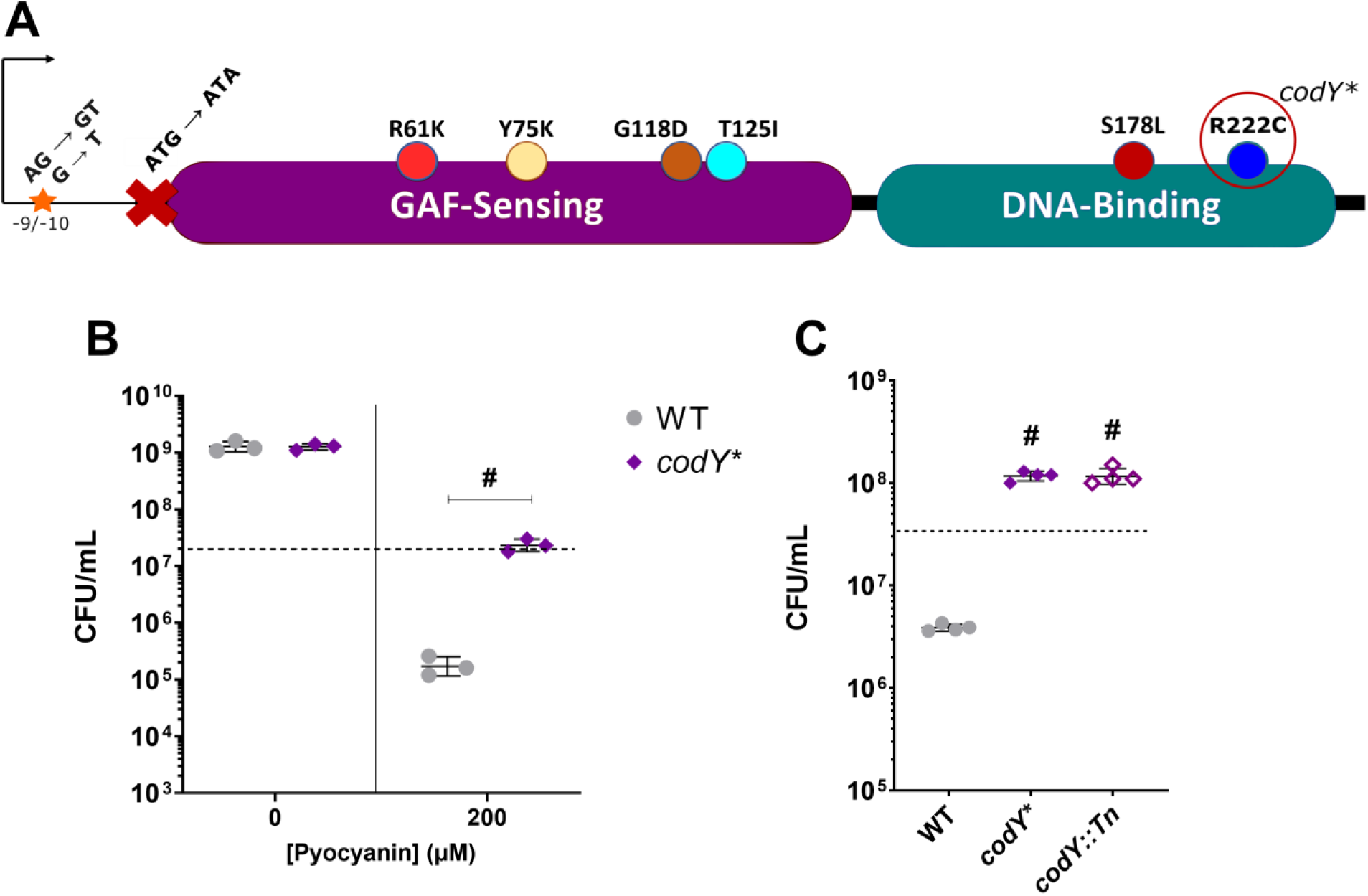
Loss of CodY function confers PYO tolerance. (**A**) Schematic of CodY-associated mutations observed in evolved isolates and where they reside within the protein and upstream sequence. The CodY^R222C^ allele is recapitulated in the *codY\** mutant. (**B**) Viable cell counts of the WT and *codY*\* mutant after 20 hours of treatment with either DMSO as a control or 200 µM PYO. Data shown are the geometric mean ± geometric standard deviation of three biological replicates. (**C**) Viable cell counts of the WT, *codY*\* mutant, and a transposon mutant of *codY* (*codY*::Tn) after 20 hours of treatment with 200 µM PYO. Data shown are the geometric mean ± geometric standard deviation of four biological replicates. (**B, C**) Dashed lines indicate the mean initial cell density (CFU/mL) for all strains at the time of PYO or DMSO addition. Significance is shown for comparison to the respective WT condition as tested by a (**B**) two-way ANOVA using Tukey’s correction or (**C**) one-way ANOVA using Dunnett’s correction for multiple comparisons (^#^*P* < 0.0001).

To determine if a CodY mutation is sufficient to recapitulate the PYO tolerance phenotype of our evolved isolates, we reconstructed the allele leading to one of the observed mutations, CodY^R222C^ (hereafter referred to as *codY\**), in the parental strain. When the *codY\** mutant was treated with 200 μM PYO, we observed significantly greater survival (∼100-fold) compared to the WT (**Fig. 2B**), while no difference was seen upon exposure to the DMSO control. In addition, a mutant with a transposon insertion in *codY* knocking out CodY activity phenocopied the *codY\** mutant when treated with PYO (**Fig. 2C**), providing further evidence that loss of CodY function mediates PYO tolerance.

We also tested whether loss of CodY function confers resistance to PYO, allowing for growth at higher PYO concentrations. We found that the *codY\** mutant exhibited greater growth than the WT at relatively low concentrations of PYO (12.5 and 25 µM), while a PYO concentration of 50 µM almost completely inhibited growth for both WT and the *codY\** mutant (**Supp. Fig. 2**). These data suggest that while a *codY\** mutation confers some resistance to low concentrations of PYO, selection of *codY* mutations was largely due to increased survival and PYO tolerance at the PYO concentrations used for experimental evolution. In addition, because we observed this moderate enhancement of PYO resistance in the *codY\** mutant (**Supp. Figs. 2B, 2C**), we tested whether a previously identified PYO resistance mutation would also engender increased PYO tolerance. It has been shown that loss of QsrR, a quinone-sensing repressor of quinone detoxification genes (44), results in PYO resistance in *S. aureus* (38), allowing for enhanced growth in up to 32 µM PYO compared to the parental strain. While our experimental conditions and growth medium are different, we observed greater sensitivity of the Δ*qsrR* mutant to 50 – 200 µM PYO compared to the WT (**Supp. Fig. 3**), indicating that although loss of QsrR allows for increased growth in the presence of lower concentrations of PYO, it does not confer increased survival or tolerance to higher concentrations of PYO in our assay conditions. Together, these data indicate that CodY loss-of-function mutations enhance *S. aureus* survival in the presence of high concentrations of PYO via a mechanism distinct from previously identified adaptive mutations.

### A CodY^R222C^ mutant exhibits expected transcriptional changes in metabolism and virulence gene expression

CodY regulates a substantial proportion of the *S. aureus* genome based on branched-chain amino acid (isoleucine, leucine, and valine) and nucleotide (GTP) availability (43, 45). In the presence of sufficient intracellular concentrations of these nutrients CodY is active and functions primarily to repress its target genes (42). As nutrients become depleted or scarce, CodY activity decreases, thereby facilitating the expression of amino acid transport and biosynthesis genes and host-targeting virulence factors. Due to the broad regulon of CodY, we determined the transcriptional response of the WT and *codY\** mutant to PYO at early (30 minutes) and late (120 minutes) time points in order to identify differentially expressed genes between the two strains that may explain the increased PYO tolerance of the *codY\** mutant (**Supp. Data File 02**). The gene expression changes in the *codY\** mutant compared to WT under control conditions at 30 minutes were consistent with previous reports of the CodY regulon (42, 43) (**Supp. Fig. 4**), indicating that CodY^R222C^ disrupts CodY-dependent regulatory activity. Among enriched pathways from overexpressed genes in the *codY\** mutant, the most prevalent were those involved in amino acid biosynthesis and metabolism, while those from downregulated genes involved responses to metal stress and protein refolding (**Supp. Fig. 4A**). Individual genes involved in these pathways were also generally among the most highly differentially expressed genes (**Supp. Fig. 4B**). We observed no significant differentially expressed genes in the *codY\** mutant compared to WT in DMSO at 120 minutes (**Supp. Data File 02**), likely due to the natural inactivation of CodY in the WT at this time point.

Overexpressed genes in the *codY\** mutant also included components of the *agr* quorum sensing system (**Supp. Fig. 4B**) that regulates virulence gene expression (46, 47). A recent report showed that *agr* activation can impart long-lived protection from oxidative stress (48). To test whether *agr* overexpression contributes to survival against PYO-mediated killing in the *codY\** mutant, we introduced an *agrA*::Tn knockout allele from the Nebraska Transposon Mutant Library (NTML) (49) into the *codY\** background. While the *agrA*::Tn mutation appeared to result in an extended lag phase as described recently (48) and seen by the lower initial cell densities in both the WT and *codY\** backgrounds (compare dashed lines for the two strains on the left and the two strains on the right in **Supp. Fig. 5**), the *codY\** mutation still provided protection comparable to the *agr*-competent strains (**Supp. Fig. 5**), suggesting that overexpression of *agr* in the *codY\** mutant is not responsible for the increased PYO tolerance.

### Induction of stress response and iron acquisition genes and suppression of metabolism characterize the response to PYO

Considering that *S. aureus* and *P. aeruginosa* are frequently found to co-infect patients, and *S. aureus* may thus be exposed to PYO, we determined the response of *S. aureus* to PYO by comparing gene expression of the WT JE2 strain in PYO to the DMSO control. To our knowledge, this is the first characterization of the *S. aureus* transcriptional response to PYO. Enriched pathways among the overexpressed genes upon PYO exposure included genes involved in iron acquisition (siderophore and heme metabolism), energy generation (menaquinone biosynthesis and the TCA cycle), and stress response to oxidative stress, metals, and DNA damage (**Fig. 3**), indicating that PYO likely generates reactive oxygen species that are known to affect metal homeostasis (50–52), causes DNA damage, and leads to metabolic alterations in *S. aureus*. Enriched pathways within the downregulated genes at the 30-minute time point were broadly comprised of amino acid transporters and diverse transcription factors (**Fig. 3A**). At the later time point, enriched pathways from downregulated genes that were independent of the effect of the natural inactivation of CodY at this time-point included pathways for amino acid and nucleic acid metabolism, energy generation, and translation (**Figure 3C** and **Supp. Fig. 6**), likely reflecting growth inhibition upon PYO exposure.

**Figure 3.**
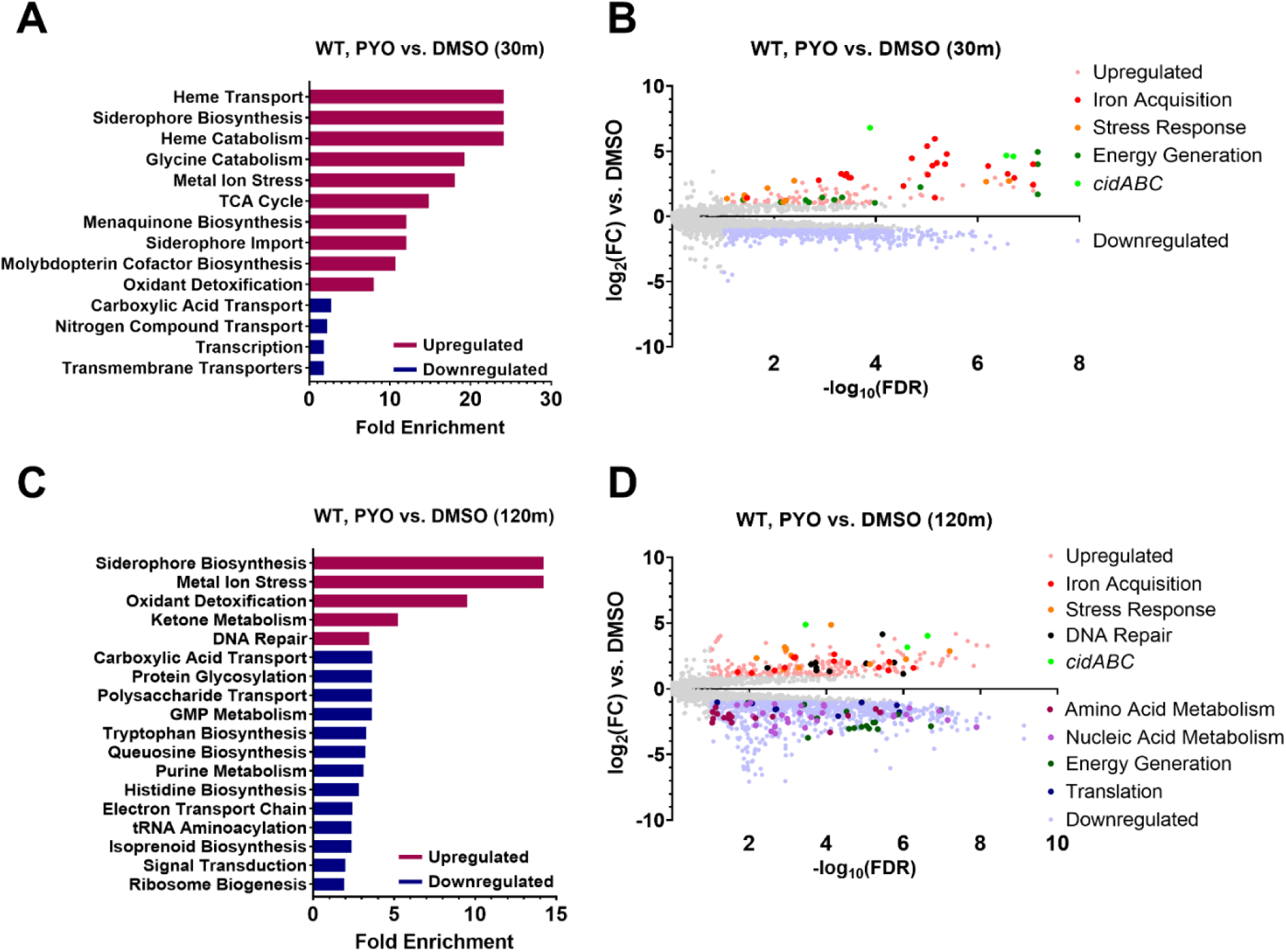
S. aureus increases expression of genes associated with iron acquisition and stress responses while suppressing metabolism- and translation-associated genes in response to PYO. Differential gene expression of WT S. aureus in response to 200 µM PYO after 30 (**A**, **B**) and 120 (**C**, **D**) minutes of incubation in PYO. (**A**, **C**) Enriched GO pathways at 30 and 120 minutes among upregulated and downregulated genes and their fold enrichment relative to the expected number of observed genes. (**B**, **D**) Volcano plots of log_2_(fold change gene expression) and -log10(false discovery rate). Upregulated genes are shown in light red and downregulated genes are shown in light blue. *Further* highlighted genes indicate the over- and under-expressed genes comprising the associated functional pathways in the legend (see **Supplementary Data File 03** for a list of the included genes). Enriched GO pathways (**C**) and differentially expressed genes (**D**) at 120 minutes include only those not also observed in the codY* mutant compared to WT from the 30-minute DMSO comparison (**Supp. Fig. 4**), but a full list can be found in **Supp. Fig. 6** and as part of Supplementary Data File 0*2*.

We also noted that *cidA*, part of the *cidABCR* operon that plays a central role in controlling carbon metabolism and programmed cell death (53, 54), was the most overexpressed gene in the WT JE2 strain at both early (∼111-fold) and late (∼29-fold) time points, while *cidB* and *cidC* were also highly upregulated (**Figs. 3B, 3D**). Considering the reported role of this operon and its individual constituents in regulating cell death (55, 56), we tested whether any of the *cid* genes were involved in PYO-dependent cell death or tolerance. We observed no effect for mutations in any of the *cidABCR* genes in either the WT or *codY\** backgrounds (**Supp. Fig. 7**), suggesting that, despite its strong overexpression, the *cid* operon does not contribute to cell death or survival in these conditions. It is possible that the strong induction of these genes is related to a metabolic or physiological shift following respiratory inhibition by PYO. Of note, the redox-sensing two-component regulator SrrAB is known to regulate *cidABC* expression (57, 58) and could be a link between PYO exposure and *cidABC* overexpression.

Since we also observed enrichment of DNA damage pathways among upregulated genes, we determined if PYO treatment led to DNA breaks using a TUNEL assay and tested whether this differentially affected the WT versus the *codY\** mutant. We observed significant DNA damage after 2-, 4-, and 20-hour treatment with PYO in the WT (**Supp. Fig. 8A**). When comparing DNA breaks in WT to the *codY\** mutant, WT exhibited a moderate, but not statistically significant increase in TUNEL staining compared to the *codY\** mutant in PYO at 2 hours, while there was no significant difference at other time points (**Supp. Figs. 8B, 8D-F**). Since we also do not observe killing following 2- and 4-hour treatment with PYO (**Supp. Fig. 8C**), these data suggest that while PYO does induce DNA breaks, these likely do not underlie the differential PYO tolerance between WT and the *codY\** mutant.

### The *codY\** mutant exhibits greater suppression of translation and expression of stress response genes than the WT

We next examined the transcriptional response of the *codY\** mutant to PYO, which was overall similar to that of the WT. However, at the early (30 min) time-point, *codY\** appeared to exhibit a more extensive repression of amino acid and nucleic acid metabolism, energy generation, and translation-associated pathways (**Supp. Figs. 9A, 9B**; contrast with **Fig. 3A, 3B**), indicating that in the *codY\** mutant PYO may be rapidly inducing a more metabolically dormant state, which has been associated with antibiotic tolerance and persistence (34, 59).

We then directly compared gene expression between the *codY\** mutant and WT in the PYO condition to identify transcriptional differences that could potentially explain the PYO tolerance of the *codY\** mutant. Apart from CodY-dependent amino acid metabolism pathways, at the early time-point, translation was enriched among the downregulated genes (**Fig. 4A** and **Supp. Fig. 10A**), consistent with the greater metabolic suppression seen in the *codY* mutant at this time-point. Further, several genes and pathways potentially involved in stress responses were upregulated in the *codY\** mutant. These included *pxpBCA* encoding 5-oxoprolinase that converts 5-oxoproline to glutamate (60), and *gltBD* encoding glutamate synthase, both of which generate glutamate that has been implicated in the response to oxidative and other stresses (61–63), and *adhC* (encoding alcohol dehydrogenase), that has been implicated in resistance to oxidative and nitrosative stress (64, 65), which were consistently upregulated at both time points (**Fig. 4A, 4B**). Additionally, the carotenoid biosynthesis pathway, and catalase-encoding *katA*, which are both important for resistance to oxidative stress (50, 66) were upregulated at the later time-point (**Fig. 4B** and **Supp. Fig. 10**) suggesting that the *codY\** mutant has a more robust stress response to PYO which may underlie its increased survival.

**Figure 4.**
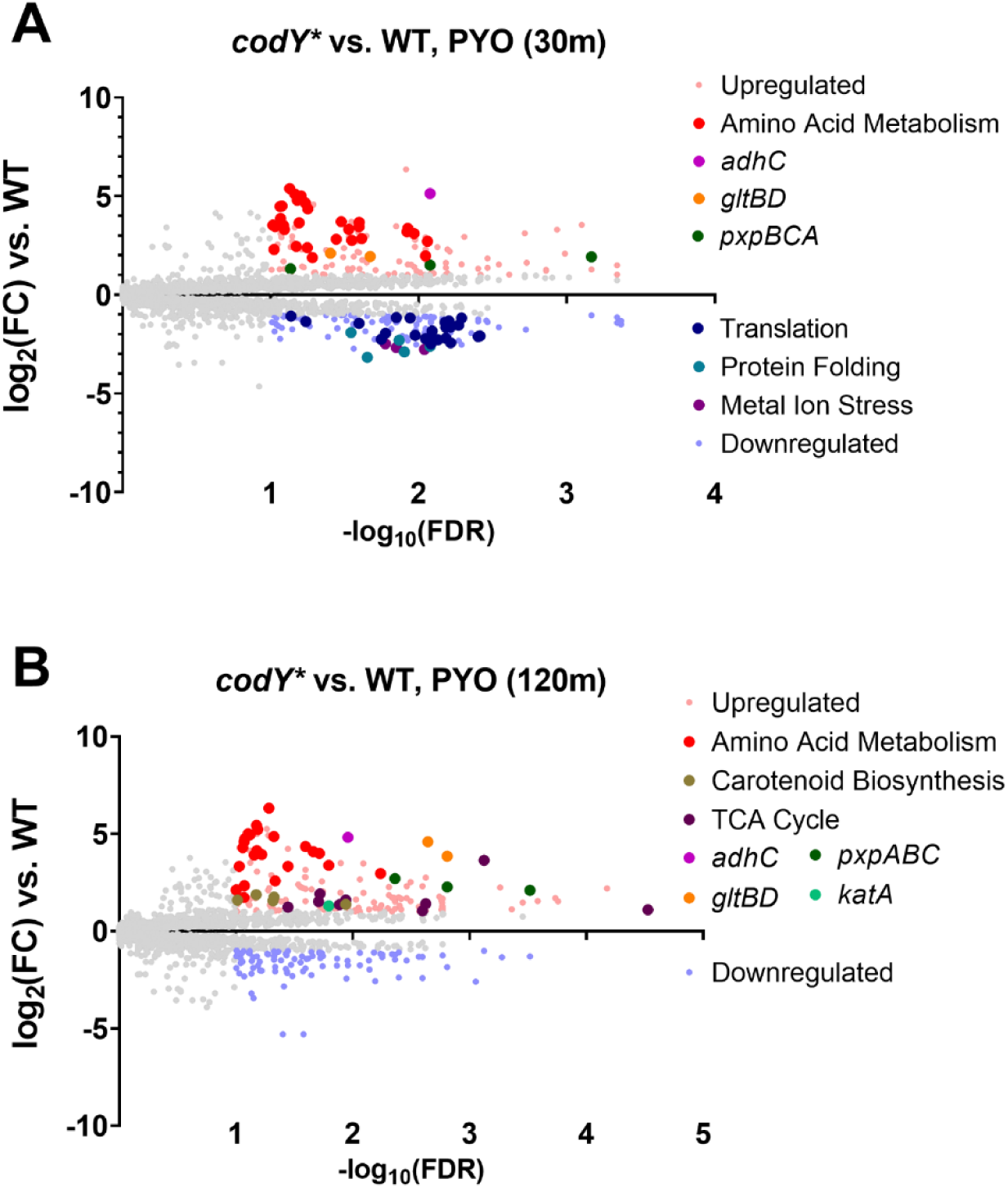
The codY* mutant exhibits enhanced expression of amino acid metabolism and stress response genes and downregulates translation compared to WT in response to PYO. Volcano plots showing log_2_(fold change gene expression) and -log10(false discovery rate) in response to 200 µM PYO in the codY* mutant compared to WT after 30 (**A**) and 120 (**B**) minutes. Highlighted genes indicate the over- and under-expressed genes comprising the associated functional pathways (see **Supplementary Data File 03** for a list of the included genes) or selected genes associated with stress responses. Enriched GO pathways for the respective volcano plots can be found in **Supp. Fig. 10**.

### ATP depletion protects *S. aureus* from the bactericidal effects of PYO

Pathways involved in ATP synthesis, nucleotide biosynthesis, translation, and respiration were enriched among downregulated genes in the *codY\** response to PYO compared to the DMSO control (**Supp. Fig. 9**), and translation-associated genes were repressed in the *codY\** PYO response compared to the WT (**Fig. 4A**). Therefore, we hypothesized that metabolic quiescence may promote protection from PYO. Additionally, ATP depletion in *S. aureus* and *E. coli* is associated with the formation of persister cells and reduced susceptibility to antibiotics (67, 68). We therefore added the proton motive force decoupling agents carbonyl cyanide m-chlorophenyl hydrazone (CCCP), a protonophore that reduces ATP synthase activity (69, 70), or sodium arsenate, which reduces ATP by forming unproductive ADP-arsenate (67, 71), to WT cultures prior to PYO addition. Addition of either 10 μM CCCP or 30 mM arsenate resulted in WT PYO tolerance comparable to that of the *codY\** mutant (**Fig. 5A**). We observed no additional protective effect of ATP depletion via CCCP addition on PYO tolerance of the *codY\** mutant (**Supp. Fig. 11**). While CCCP did not interfere with growth in the absence of PYO, arsenate arrested growth at the point of addition.

**Figure 5.**
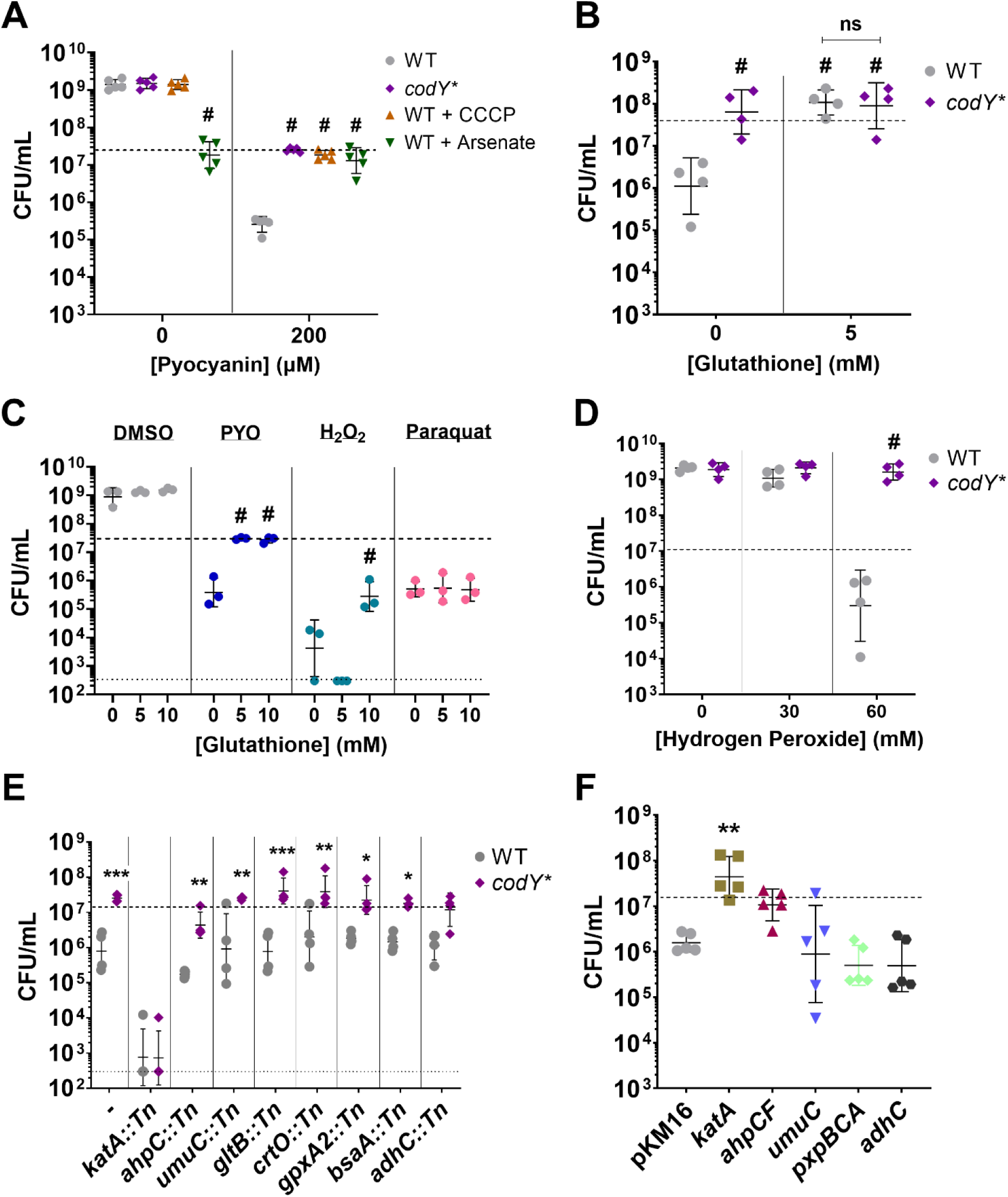
Metabolic suppression and hydrogen peroxide stress response mediate PYO tolerance. Viable cell counts of the indicated *S. aureus* strains after 20-hour treatment with the stated reagents. (**A**) Tolerance to 200 µM PYO of the *S. aureus codY\** mutant, and WT in the presence of the ATP depleting agents CCCP (10 µM) and sodium arsenate (30 mM). (**B**) Tolerance to 200 µM PYO of the WT and the *codY\** mutant in the presence of glutathione or sterile water as a control. (**C**) Effect of glutathione on WT survival during treatment with 200 µM PYO, 60 mM hydrogen peroxide, or 0.1 mM paraquat. (**D**) Susceptibility of WT and the *codY\** mutant to hydrogen peroxide. (**E**) PYO sensitivity of transposon mutants disrupting overexpressed genes from RNA-seq in WT and the *codY\** mutant backgrounds. (**F**) Overexpression of selected genes with their native promoters from a multi-copy plasmid. pKM16 expresses the fluorescent protein dsRed3.T3 from the *sarA1* promoter of *S. aureus* and is used as a control. Supplemental reagents were added at the indicated concentrations immediately prior to addition of 200 µM PYO, DMSO, or sterile water. Dashed lines indicate the mean initial cell density (CFU/mL) for all strains prior to addition of PYO, DMSO, or other reagents, while the lower dotted lines in (**C**) and (**E**) indicate the limit of detection. Data shown are the geometric mean ± geometric standard deviation of the following numbers of biological replicates: (**A**) five, (**B**) four, (**C**) three, (**D**) four, (**E**) four, (**F**) five. Significance is shown relative to (**A**) the respective WT condition, (**B**) WT without glutathione, or the indicated strains, (**C**) the respective no glutathione condition, (**D**) the respective WT condition, (**E**) the respective WT mutant, or (**F**) pKM16, and was determined by (**A-E**) a two-way ANOVA using (**A**) Dunnett’s, (**B**, **C**, **D**) Tukey’s, or (**E**) Šídák’s correction, or (**F**) one-way ANOVA using Dunnett’s correction for multiple comparisons (\**P* < 0.05, \*\**P* < 0.01, \*\*\**P* < 0.001, ^#^*P* < 0.0001).

Despite their metabolic dormancy, persister cells maintain some active metabolic processes (72, 73) which can be necessary for the protective effects of persistence (74–76). To determine whether CCCP-mediated protection from PYO still required active metabolism, we tested the effect of higher CCCP concentrations on both the WT and *codY\** mutant strains. While 40 μM CCCP did not adversely affect growth in the DMSO control, this concentration resulted in an increased susceptibility to PYO in both the WT and *codY\** mutant strains (**Supp. Fig. 11**), suggesting that active metabolism is likely required for basal PYO tolerance in the WT and increased PYO tolerance in the *codY\** mutant. Together, these data indicate that partial metabolic suppression in the *codY\** mutant may contribute to PYO tolerance.

### Mitigation of hydrogen peroxide toxicity is protective against PYO

PYO toxicity is in part mediated by the generation of diverse ROS (31, 77), and the oxidative stress response pathway is induced in both the WT and *codY\** mutant upon PYO exposure (**Fig. 3** and **Supp. Fig. 9**). To determine whether the PYO-induced cell death observed in the WT was mediated by ROS, we screened *S. aureus* mutants deficient in different ROS responses for sensitivity to PYO. These included mutants in the catalase (*katA*) and alkyl hydroperoxide reductase (*ahpC, ahpF*) hydrogen peroxide detoxification systems (78, 79), the peroxiredoxins thiol peroxidase (*tpx*) and thiol-dependent peroxidases (*bsaA, gpxA2*) predicted to detoxify peroxides (80), both superoxide dismutases (*sodA, sodM*) (81), and a biosynthesis gene for bacillithiol (*bshA*) predicted to function in redox homeostasis (80). Mutations in *katA*, *ahpC*, and *ahpF* sensitized WT to PYO-mediated killing (**Supp. Fig. 12**), indicating that detoxification of hydrogen peroxide is critical for PYO tolerance. Interestingly, a mutant of *perR*, a master repressor of the oxidative stress response (50, 51), phenocopied the PYO tolerance of the *codY\** mutant (**Supp. Fig. 12**). A *perR* mutant constitutively expresses several stress response genes, including *katA*, *ahpCF*, *dps*, *trxB*, and *ftnA* (50), corroborating that an enhanced ROS response is protective against PYO.

To further test whether ROS were responsible for the killing effect of PYO, we supplemented WT with glutathione, a well-known antioxidant (82–84), and observed substantially increased survival of the WT, comparable to the *codY\** mutant (**Fig. 5B**). Similar to the CCCP treatment, we observed no additional protective effect of glutathione on the *codY\** mutant. Glutathione supplementation also provided partial protection against hydrogen peroxide, but not paraquat-induced superoxide (**Fig. 5C**), indicating that hydrogen peroxide is a primary mediator of cell death under these conditions and consistent with the enhanced PYO sensitivity of the *katA* and *ahpCF* mutants (**Supp. Fig. 12**).

In the *codY\** mutant, *katA* was overexpressed 2.2- to 2.5-fold compared to WT in DMSO (**Supp. Data File 02**) and in the later response to PYO (**Fig. 4B**), suggesting that the *codY\** mutant exhibits an elevated basal tolerance to hydrogen peroxide and a greater response to ROS stress compared to WT. We found that the *codY\** mutant exhibited increased resistance to hydrogen peroxide (**Fig. 5D**) but not superoxide (**Supp. Fig. 13**). Together, these data indicate that PYO-mediated toxicity is primarily mediated by hydrogen peroxide and that *codY\** confers greater tolerance to this ROS.

Given the increased tolerance of the *codY\** mutant to ROS and specifically PYO-generated ROS, and the apparent requirement of active metabolism for this, we selected several stress response related genes whose transcripts were elevated in the *codY\** mutant either upon PYO exposure or compared to WT in response to PYO to assess their role in PYO tolerance of the *codY\** mutant. In addition to *katA*, *ahpC*, *bsaA*, and *gpxA2*, we selected the biosynthesis pathway of carotenoids (*crtO*) which function to alleviate hydrogen peroxide stress (66, 85); alcohol dehydrogenase (*adhC*) which is part of the Rex regulon responsive to redox stress and oxygen limitation (86, 87); MucB (*umuC*), an error-prone DNA polymerase which functions in DNA repair (88); and glutamate synthase (*gltB*) since glutamate utilization can mediate protection against oxidative stress (61). We transduced transposon mutations in each of these genes to the *codY\** mutant background. When challenged with PYO, we found that loss of *katA* (*codY* katA*::Tn) substantially reduced PYO tolerance and abolished the protective effects of *codY\**, suggesting that *katA* is required for PYO tolerance in the *codY\** mutant (**Fig. 5E**). In contrast, while the *codY* ahpC*::Tn strain showed lower survival compared to the *codY\** mutant, it still showed higher tolerance compared to the *ahpC*::Tn strain.

Finally, we asked whether overexpression of these or other stress-related genes would be sufficient to confer PYO tolerance. When overexpressed from a multi-copy plasmid, we observed that *katA* clearly protected WT from PYO-mediated killing at levels similar to *codY\** and expression of *ahpCF* exhibited a moderate, though not statistically significant, protective effect (**Fig. 5F**). In contrast, overexpression of *umuC*, *pxpBCA*, or *adhC* was not sufficient to protect WT from PYO-mediated killing (**Fig. 5F**). Taken together, these results indicate that an enhanced response to hydrogen peroxide stress is sufficient to mediate PYO tolerance and that the overexpression of *katA* in the *codY\** mutant contributes to its increased survival in PYO.

### Experimentally evolved mutations in CodY are present in genomes of *S. aureus* clinical isolates

In our experimental evolution, we identified *codY* mutations in each of 18 sequenced isolates from two independently evolved populations. In total, we observed nine different mutations: seven in the coding sequence, an ablation of the start site, and two promoter mutations. Given that we didn’t isolate the same coding sequence mutation from both independently evolved populations, and that these mutations probably led to loss of CodY function, there is likely a large mutational space of CodY-inactivating alleles that may be selected for upon exposure to redox stress. Further, it has been shown that a *codY* deletion leads to increased *in vivo* virulence in a USA300 strain (89), suggesting that this could be an additional selective pressure for *codY* inactivating alleles.

To test whether *codY* mutations are seen in publicly available genomes of *S. aureus* strains, we queried the JE2 CodY protein sequence against a set of 63,983 *S. aureus* genomes from the NCBI Pathogen Detection Database (**Supp. Data File 04**). Interestingly, we identified multiple isolates which had CodY mutations that we observed in our study, including T125I, S178L, and R222C, as well as a variety of other mutations throughout the protein, suggesting that these mutations can and do arise in natural isolates (**Supp. Data File 04**).

## DISCUSSION

Constituents of polymicrobial communities can exhibit competitive behaviors that affect other community members (9). Such selective pressures within these communities can promote adaptations that maintain or shift the balance of community form and function (4), but the breadth of these mechanisms is not well-characterized. In this study, we use experimental evolution to investigate the adaptive response of *S. aureus*, a widespread pathogen frequently identified in antibiotic-resistant and polymicrobial infections, to the redox-active *P. aeruginosa* antimicrobial, PYO. We show that recurrent treatment with a bactericidal concentration of PYO selects for increased *S. aureus* survival mediated by loss-of-function mutations in the pleiotropic transcriptional repressor, CodY. Transcriptional analysis during PYO treatment indicates that the *codY\** mutant shows a stronger repression of translation-associated genes and greater expression of certain stress response genes compared to WT, suggesting that transcriptional changes in the *codY\** mutant confer PYO tolerance. Consistent with this hypothesis, we observed that, individually, metabolic suppression or overexpression of catalase was sufficient to impart PYO tolerance to the WT. Our results suggest a multifaceted adaptive response to antimicrobial-induced reactive oxygen stress that reduces lethal cellular damage through reduced metabolism and enhanced ROS detoxification.

PYO and other phenazines have diverse functions in *P. aeruginosa* physiology (90–94), pathogenesis (30, 95), and interbacterial competition (36, 37). The ability of PYO to accept and donate electrons enables it to interfere with respiratory processes of other species (36, 40, 96) and generate toxic ROS through the reduction of molecular oxygen (31, 37). Although PYO is frequently undetectable in CF sputum even during colonization with *P. aeruginosa* (97–99), PYO production has been observed during human disease, including in ear and CF lung infections (28, 29) and several lines of evidence suggest a potential role in infection. Culture in *ex vivo* CF sputum (100), *in vitro* in CF-sputum mimicking medium (101) or in the presence of anaerobic products frequently found in CF lung environments or breath condensates (102–105) can induce expression of PYO biosynthesis genes or PYO production. In addition, overproduction of PYO (106) and regulatory rewiring that maintains PYO production (107) have been observed in CF isolates, and one study showed an association between increased PYO production and isolates from pulmonary exacerbation sputum samples (108).

Recently published studies have used experimental evolution to identify diverse adaptive mechanisms of *S. aureus* to *P. aeruginosa* antagonism. Loss-of-function mutations in the aspartate transporter, *gltT,* led to increased survival of *S. aureus* during surface-based co-culture competition with *P. aeruginosa* (109), as selective pressure under those conditions was primarily related to competition for amino acids. While we observe downregulation of *gltT* and the glutamate transporter *gltS* in response to PYO in both WT and the *codY\** mutant (**Supp. Data File 02**), we would not expect similar selective pressures in our assay. In a separate study, *S. aureus* evolved in the presence of *P. aeruginosa* supernatant showed strain-dependent acquisition of resistance that converged on staphyloxanthin (carotenoid) production and the formation of small colony variants (SCVs) (110). Interestingly, in two different strains, 1 out of 5 populations each encoded a mutation in *codY*, one being intergenic and the other a non-synonymous mutation. While we also observed overexpression of staphyloxanthin biosynthesis genes (**Fig. 4B**), we did not observe increased sensitivity to PYO in a *crtO* knockout mutant (**Fig. 5E**). It is likely that differences in the primary phenotype (resistance versus tolerance) and the mixture of inhibitory factors present in *P. aeruginosa* supernatant can explain this difference, but also suggests that the consequences of *codY* mutation can facilitate protective phenotypes in other conditions.

Selection of *S. aureus* mutants that can grow in the presence of PYO identified SCVs and mutations in *qsrR* as PYO resistance determinants in previous studies (38, 96). The antimicrobial activity of PYO is considered to involve two functions: ETC inhibition and generation of ROS (31, 32). SCVs likely evade ETC inhibition and ROS generation due to reduced respiration. Alternatively, *qsrR*-mediated responses are reported to detoxify PYO leading to PYO resistance (38). It seems likely that ETC inhibition was critical for the bacteriostatic effects of PYO and subsequent PYO resistance mechanisms in these studies. In our conditions, we observed substantial induction of stress response transcripts (**Fig. 3**), suggesting that ROS generation is a major effect of PYO. In particular, expression of *katA*, *ahpCF,* and the cytoplasmic iron-sequestering protein *dps* that protects DNA from hydrogen peroxide (111, 112) were highly overexpressed in response to PYO in both the WT and *codY\** mutant (**Fig 3, Supp. Fig. 9**, and **Supp. Data File 02**). Additionally, genes involved in distinct iron acquisition systems were among the most upregulated genes in response to PYO (**Fig. 3** and **Supp. Fig. 9**). In *S. aureus*, metal acquisition and homeostasis genes are integrated into the regulons of peroxide stress response regulators such as PerR and Fur (50, 51), likely due to the iron-dependent functions of redox proteins such as catalase. Hydrogen peroxide can also induce expression of heme and iron uptake genes (52). Together with the observation that overexpression of *katA* is sufficient to induce PYO tolerance (**Fig. 5F**), these data suggest that the bactericidal effects of PYO in *S. aureus* are primarily driven by the generation of peroxides. Interestingly, this is in contrast to observations in *A. tumefaciens* where superoxide dismutase was the critical ROS stress protein mediating PYO tolerance (37). It is possible that this difference reflects differences between the physiology of the two organisms and the experimental conditions.

CodY regulates approximately 5-28% of the *S. aureus* genome depending on the strain and experimental conditions (43, 45), predominantly by repressing genes that function in metabolism and virulence factor production (41, 42). Therefore, the presence of mutations associated with *codY* raised the immediate possibility that transcriptional changes prior to or during PYO treatment contribute to PYO tolerance. Indeed, we observed differential expression of multiple stress response genes during PYO treatment in the *codY\** mutant compared to WT (**Fig. 4B, Supp. Fig. 4B**, and **Supp. Data File 02**). Most of these genes (*katA*, *crtO*, *gpxA2*, *gltB*) are known or predicted to detoxify peroxides. However, among these overexpressed genes, only loss of *katA* fully sensitized the *codY\** mutant to PYO (**Fig. 5E**). Overexpression of *katA* was also observed in the *codY\** mutant in the absence of PYO (**Supp. Data File 02**), likely contributing to its enhanced resistance to hydrogen peroxide (**Fig. 5D**) but not superoxide (**Supp. Fig. 13**). Based on the protection provided by the *codY\** mutation even in an *ahpC* mutant (**Fig. 5E**), it is likely that most of the toxic effects of PYO are borne from high levels of hydrogen peroxide that are more effectively detoxified by the functionally intact catalase in this mutant (79). Based on our results, the enhanced stress response could contribute to the increased virulence of a CodY mutant (89) alongside the de-repression of virulence factors by conferring protection from host defenses (42, 43).

In addition to overexpression of stress responses, the *codY\** mutant also exhibited a strong, early reduction of translation-associated gene expression compared to the WT (**Fig. 4A**), and several amino acid and nucleotide biosynthesis pathways as well as the electron transport chain were all enriched among genes downregulated early in the *codY\** mutant upon PYO exposure (**Supp. Fig. 9**). This indicated a rapid metabolic suppression in the *codY\** mutant by PYO and artificially depleting ATP was sufficient to protect WT from PYO-mediated killing (**Fig. 5A**). Consistent with lowered metabolism leading to PYO tolerance, it has been previously shown that the development of antibiotic tolerance and persistence is mediated by metabolic restriction, where reduced translation, ATP levels, or growth rate can enhance survival against antimicrobials (67, 113). In this context, suppressing metabolism may also contribute to protection by reducing the accumulation of PYO-generated ROS. Combined with increased expression of *katA*, our results suggest that both responses contribute to PYO tolerance via overlapping mechanisms.

Notably, in our experiments, although a *perR* mutant showed high PYO tolerance (**Supp. Fig. 12**), and we saw induction of many genes within its regulon upon PYO exposure, *perR* mutations were not present in any of our evolved isolates. Similarly, we did not identify any promoter mutations in *katA* that would lead to overexpression. This could be due to the small number of populations we evolved, minor fitness costs associated with *katA* promoter mutations and *perR* loss of function mutations, or unique selective pressures exerted by our experimental evolution protocol. Alternatively, it is possible that the levels of *katA* induction accessible to such specific mutations are lower than what we see via plasmid overexpression, and not sufficient by itself to lead to high levels of PYO tolerance. Instead, the experimental evolution selected for *codY* mutations that led not only to *katA* overexpression, but also metabolic suppression, and the combination of both these effects likely led to the observed high PYO tolerance.

It has been suggested that mutations in regulators can facilitate adaptation by optimizing regulatory processes toward a new niche, via increasing expression of critical biochemical capabilities, or inhibiting wasteful or damaging metabolic processes (114). Here we show that mutations in global regulators may also allow access to unique peaks in the fitness landscape due to pleiotropic effects on cellular metabolism, thereby facilitating multiple distinct protective responses.

## MATERIALS AND METHODS

### Bacterial strains and growth conditions

All strains and plasmids used in this study are described in **Supplemental Table 2**. Bacteria were cultured at 37 °C with shaking at 300 rpm in modified M63 medium (13.6 g/L, KH_2_PO_4_, 2 g/L, (NH_4_)_2_SO_4_, 0.4 µM ferric citrate, 1 mM MgSO_4_; pH adjusted to 7.0 with KOH) supplemented with 0.3% glucose, 1x ACGU solution (Teknova), 1x Supplement EZ (Teknova), 0.1 ng/L biotin, and 2 ng/L nicotinamide for all experiments. Luria-Bertani (LB) broth (KD Medical) or LB agar (Difco) was used for cloning and enumerating CFU and supplemented with 10 µg/mL erythromycin, 10 µg/mL chloramphenicol, or 100 µg/mL carbenicillin as required.

Pyocyanin (from *Pseudomonas aeruginosa*, Sigma-Aldrich [P0046]) (PYO) was resuspended to a concentration of 10 mM in dimethyl sulfoxide (DMSO), stored at -30°C, and used at the concentration(s) indicated for each experiment.

### Pyocyanin tolerance assays

Overnight cells were washed once in sterile PBS and normalized by their OD_600_. Washed cells were then inoculated to an OD_600_ of 0.1 in 1 mL of fresh, pre-warmed M63 and incubated for 2 hours in 14 mL polystyrene test tubes. After incubation, PYO at the indicated concentration (or the same volume of DMSO as a control) was added and the cells were incubated for an additional 20 hours. CFUs were enumerated by spot plating in triplicate. PYO tolerance assays shown in **Fig. 1D** and **Supp. Fig. 1** were performed as described below. For experiments using hydrogen peroxide (Sigma-Aldrich [H1009]), paraquat (Fisher Scientific [US-PST-740]), carbonyl cyanide m-chlorophenyl hydrazone (CCCP; Sigma-Aldrich [215911-250MG]), or sodium arsenate (Sigma-Aldrich [S9663-50G]), the indicated supplement was added immediately prior to the addition of PYO.

### Laboratory evolution of PYO tolerance

Two independent populations of *S. aureus* JE2 were grown overnight in M63, washed in PBS, and inoculated to an OD_600_ of 0.05 into 125 mL Erlenmeyer flasks containing 5 mL of fresh, pre-warmed M63 and grown to an OD_600_ of ∼0.25. 1 mL of culture was transferred to a sterile 14 mL test tube containing 20 μL of 10 mM PYO – yielding a final concentration of 200 μM PYO – and incubated for a further 20 hours. Following incubation, 20 μL of culture was removed to enumerate CFU and the remaining culture pelleted by centrifugation. Pelleted cells were resuspended in fresh M63 and allowed to recover overnight in the absence of PYO. Overnight recovered cells were then diluted as above, and the procedure repeated up to six times. Populations after treatment 5 for Population A and treatment 7 for Population B were streaked out on LB plates, and individual colonies were selected from these to test for PYO tolerance, and for whole-genome sequencing.

The PYO tolerance assays shown in **Fig. 1D** and **Supp. Fig. 1** were similarly performed for the indicated isolates, by enumerating CFU after 20 hours of PYO treatment.

### Construction of mutant strains

Primers used for cloning and verification are described in **Supplemental Table 3**. The location of transposon insertions in strains acquired from the Nebraska Transposon Mutant Library (NTML) were verified by PCR and Sanger sequencing. Transductions were performed using φ11 or φ80 based on previously described procedures (115). Briefly, overnight cultures bearing the transposon of interest were diluted 1:100 in a 125 mL Erlenmeyer flask containing 10 mL of BHI and grown for 2 hours (OD_600_ ∼1.0) at 37°C with shaking at 300 rpm. Then, cultures were supplemented with 150 µL of 1M CaCl_2_ followed by the addition of 1-10 µL of empty phage, and incubation was continued overnight. Culture lysates were centrifuged for 10 minutes at ∼4,000 rcf and the supernatant filtered with a 0.45 µm syringe filter. Subsequently, 100 to 1000 µL of phage lysate was used to transduce 1 mL of overnight culture of the recipient strain supplemented with 15 µL of 1M CaCl_2_ in 15 mL conical tubes. Cultures were incubated as above for 20 minutes, at which point 200 µL of 200 µM sodium citrate was added, mixed by inversion, and centrifuged as above. Cells were then resuspended in BHI containing 1.7 mM sodium citrate, incubated as above for 1 hour, and then centrifuged as above. Transduced cells were concentrated 2-fold and 100-200 µL plated on at least three BHA plates containing 10 µg/mL erythromycin and 1.7 mM sodium citrate. Cells were allowed to grow at 37°C for up to 48 hours.

Cloning for site-directed mutagenesis was performed using pIMAY* largely as previously described (116). Briefly, ∼1 ug of purified plasmid isolated from *E. coli* IM08B was transformed into electrocompetent *S. aureus* JE2 and directly selected for integration by incubation at 37°C as previously described (117). Integration was confirmed following additional overnight culture under antibiotic selection at 37°C using primers specific for plasmid integration. Integrated strains were then cultured at 28°C overnight without selection and plated on LB agar containing 20 mM para-chlorophenylalanine (PCPA) and 50 ng/mL anhydrotetracycline grown at 28°C. Colonies were screened by PCR and Sanger sequencing to identify the mutant allele.

### Pyocyanin growth curves

Overnight cells were washed once in sterile PBS, OD normalized, and inoculated to an OD_600_ of ∼0.05 in fresh, pre-warmed M63. 100 μL of culture was aliquoted in duplicate to the wells of a 96-well plate. Plates were parafilmed to reduce evaporation and cells were grown for 20 hours with shaking (807 cpm) at 37°C in a BioTek Synergy H1 microplate reader (Agilent). Optical density values are adjusted by the background value at T0.

### Quantifying DNA damage using a TUNEL assay

Overnight cultures were washed and normalized as described above for the PYO tolerance assays. Washed cells were inoculated to a calculated OD_600_ of 0.1 into 125 mL Erlenmeyer flasks containing 12 mL of fresh, pre-warmed M63 and incubated at 37°C for 2 hours with shaking at 300 rpm. After incubation, PYO at a final concentration of 200 µM or an equal volume of DMSO as a control was added and flasks were returned to the incubator. At the indicated time points, cells were pelleted by centrifugation at 14,400 rcf for 5 minutes, washed once with ice-cold PBS, and fixed on ice for 30 minutes in 1.5 mL of a 2.66% solution of PBS-paraformaldehyde (PFA). After initial fixation, cells were pelleted by centrifugation at 20,000 rcf for 3 minutes and washed with PBS to remove residual PFA. Washed cells were then resuspended in 1.25 mL of ice-cold 56% ethanol and stored for at least 24 hours at -30°C prior to TUNEL staining. TUNEL staining was performed using the APO-DIRECT Kits (BD Biosciences [51-6536AK, 51-6536BK]) according to the manufacturer’s instructions. Stained cells were analyzed using an Apogee MicroPLUS flow cytometer (ApogeeFlow Systems Inc). *S. aureus* cells were gated using medium and large angle light scatter. Fluorescently labeled DNA was excited using a 488-nm laser and collected using 515 and 610 emission filters for FITC and propidium iodide, respectively, and analyzed using FlowJo (v10.1). Comparisons were made using the Overton method to identify the proportion of the population that exhibits fluorescence compared to the control condition.

### Whole-genome sequencing

Genomic DNA from *S. aureus* was isolated using a DNeasy Blood & Tissue Kit (Qiagen) with the addition of 5 μg/mL lysostaphin (Sigma) to the pretreatment regimen described for gram-positive bacteria. Sequencing libraries were prepared using the Illumina Nextera XT DNA Library Preparation Kit according to the manufacturer’s instructions and sequenced by the CCR Genomics Core using a NextSeq 550 75 Cycle High Output kit for single-end sequencing, or the 150 Cycle Mid Output kit or High Output kit for paired-end sequencing. Genomes were assembled with breseq 0.33.1 (118) using the JE2 reference genome on NCBI (NZ_CP020619.1) as a reference.

### RNA-sequencing and analysis

Cells were cultivated in 125 mL Erlenmeyer flasks containing 10 mL of fresh, pre-warmed M63 medium at 37°C with shaking at 300 rpm. After the indicated treatment time bacterial RNA was stabilized using RNAprotect Bacteria Reagent (Qiagen) according to the manufacturer’s instructions and stored at -80°C prior to RNA isolation. RNA was isolated using a Total RNA Plus

Purification Kit (Norgen) with some modifications for *S. aureus*. Briefly, cryo-preserved bacterial pellets were resuspended in 100 μL of TE buffer containing 3 mg/mL lysozyme and 50 μg/mL lysostaphin and incubated for 30 minutes at 37°C. Volumes of Buffer RL and 95% ethanol used in the protocol were increased to 350 μL and 220 μL, respectively. Following elution of total RNA, any remaining genomic DNA was removed by TURBO DNase (Thermo Fisher) treatment using the two-step incubation method as detailed in the manufacturer’s instructions. Removal of genomic DNA was confirmed by PCR.

Ribosomal RNA was then removed using Ribo-Zero rRNA Removal Kit for gram-positive Bacteria (Illumina) according to the manufacturer’s instructions. Removal of rRNA was confirmed by electrophoresis using an Agilent TapeStation. RNA libraries were prepared using NEBNext Ultra II Directional RNA Library Prep Kit for Illumina (New England BioLabs) and sequenced using a NextSeq 550 75 Cycle High Output kit for single-end sequencing.

Sequence files were pre-processed using fastp (119). Alignment was performed using Kallisto (120) and analyzed using EdgeR (121) and RStudio. Two independent RNA-seq experiments were performed and the replicate alignments were combined for analysis. Differential gene expression between conditions was performed with the glmQLFit function in EdgeR using an FDR significance of < 0.1 and a log_2_-fold cutoff of ≥ 1 or ≤ -1. Enriched pathways were identified using Gene Ontology Resource (www.geneontology.org) by the PANTHER Overrepresentation Test (released 20240226) (Annotation Version and Release Date: GO Ontology database DOI: 10.5281/zenodo.10536401 Released 2024-01-17) after conversion of differentially expressed JE2 locus tags to NCTC8325-4 using *Aureo*Wiki (122). Pathway enrichment was tested using Fisher’s exact test corrected for the false discovery rate. All scripts used to generate results and run the above programs are provided in **Supplemental Data File 05**. Processed data files for each replicate are available in **Supplemental Data File 06**.

### Statistics

Statistical analysis of data was performed using GraphPad Prism 9 (GraphPad Software, San Diego, CA, United States). Significance was determined by one-way or two-way analysis of variance (ANOVA) as indicated in the figure legends. Log-scale values were log-transformed prior to statistical analysis.

### Data Availability

The whole genome and RNA sequencing data have both been deposited at NCBI Short Read Archive (SRA) associated with BioProject PRJNA1122578.

## ACKNOWLEDGMENTS

We would like to acknowledge the Center for Cancer Research (CCR) Genomics Core for RNA-sequencing and whole-genome sequencing, and the Brinsmade lab (Georgetown University) for providing the pKM16 plasmid. We thank Susan Gottesman, Gisela Storz, Tiffany Zarrella, Kalinga Pavan Thushara Silva, and Stefan Katharios-Lanwermeyer for comments on the manuscript, and members of the Gottesman, Ramamurthi, and Khare labs for discussion and feedback throughout the study. This work used the computational resources of the NIH High Performance Computing Biowulf Cluster (http://hpc.nih.gov). This work was supported by the Intramural Research Program of the NIH, National Cancer Institute, Center for Cancer Research.

