## Supplemental Figures and Supplemental Tables for "Mutations in the *Staphylococcus aureus* Global Regulator CodY Confer Tolerance to an Interspecies Redox-Active Antimicrobial"

### SUPPLEMENTARY FIGURES

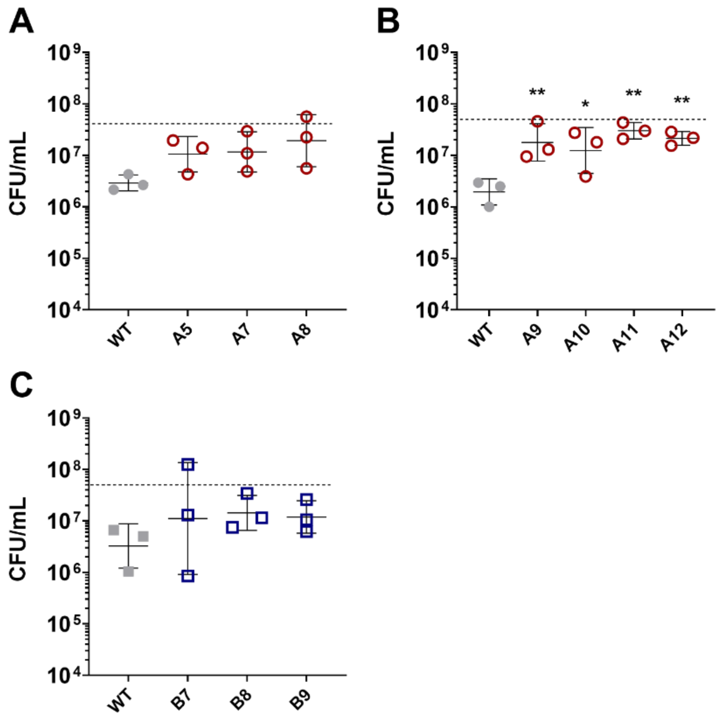

**Supplemental Figure 1. PYO tolerance of experimental evolution isolates**. PYO tolerance of terminal isolates from population A (**A, B**) and population B (**C**). Values indicate *S. aureus* viable cell counts after 20 hours of treatment with 200 μM PYO. The dashed lines indicate the mean initial cell density (CFU/mL) at the time of PYO addition. Data shown are the geometric mean ± geometric standard deviation of three biological replicates. Significance is indicated for comparison to the parent strain as determined by a one-way ANOVA using Dunnett's correction for multiple comparisons. ($^{*}P$ < 0.05, $^{**}P$ < 0.01).

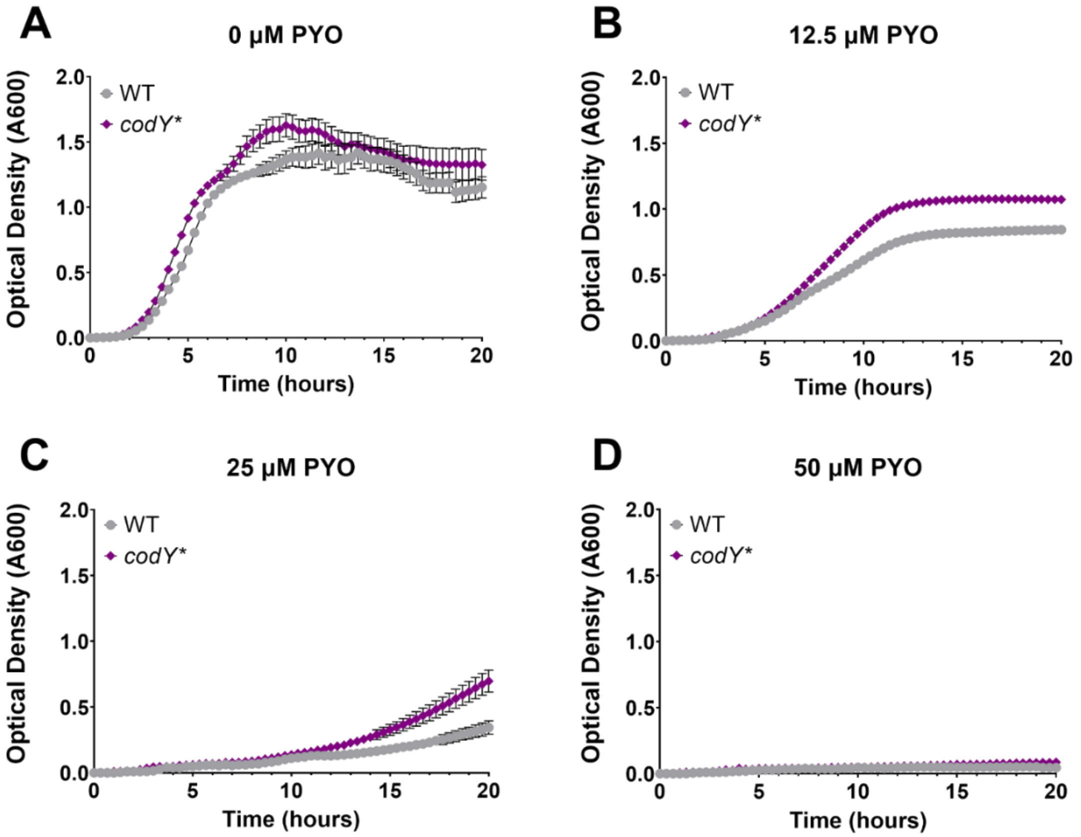

**Supplemental Figure 2. Growth of the *codY** mutant in low concentrations of PYO is modestly greater than WT**. Growth curves shown as $OD_{600}$ measurements of the WT and the *codY** mutant in M63 containing (**A**) 0 µM PYO (DMSO control), (**B**) 12.5 µM PYO, (**C**) 25 µM PYO, and (**D**) 50 µM PYO. Data shown are the mean ± standard error of seven biological replicates.

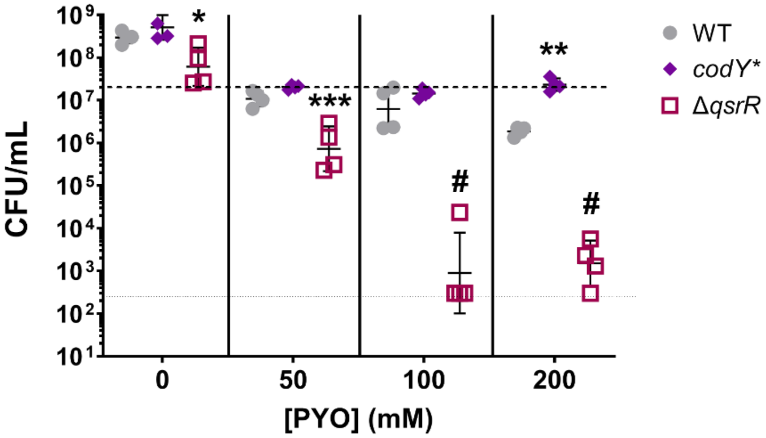

**Supplemental Figure 3. A Δ*qsrR* mutant exhibits reduced tolerance to PYO**. Viable cell counts are shown for the WT, and the *codY** and Δ*qsrR* mutants after 20-hour treatment with the indicated concentration of PYO. Data shown are the geometric mean ± geometric standard deviation of four biological replicates. The upper, dashed line indicates the mean initial cell density (CFU/mL) for all strains and the lower, dotted line indicates the limit of detection. Significance is shown for comparisons to the respective WT condition, as determined by a two-way ANOVA using Dunnett's correction for multiple comparisons. ($^{*}P < 0.05$, $^{**}P < 0.01$, $^{***}P < 0.001$, $^{\#}P < 0.0001$).

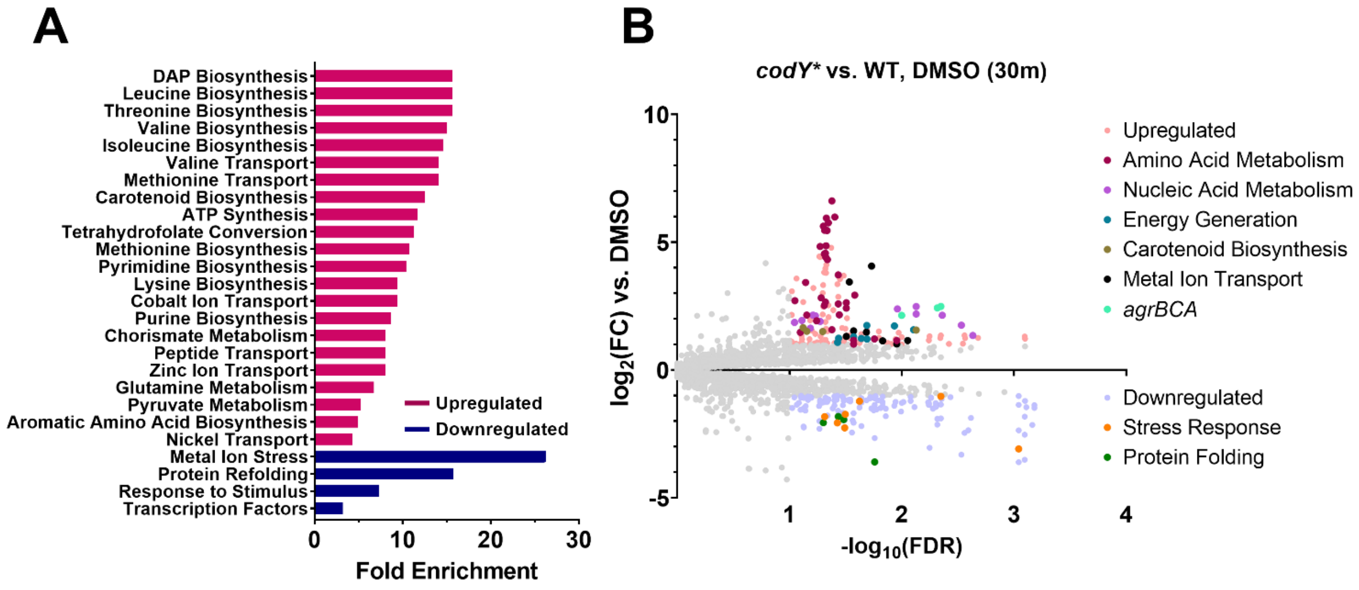

**Supplemental Figure 4. The transcriptional profile of the *codY** mutant compared to WT is consistent with loss of CodY activity**. Differential gene expression of *codY** compared to WT after 30 minutes of incubation in DMSO. (**A**) Enriched GO pathways from upregulated and downregulated differentially expressed genes. (**B**) Volcano plot of $log_2$(fold change gene expression) and -log10(false discovery rate). Upregulated genes are shown in light red and downregulated genes are shown in light blue. Individual genes from several pathways in (**A**) are further highlighted. A list of genes included in each pathway can be found in **Supplementary Data File 03**.

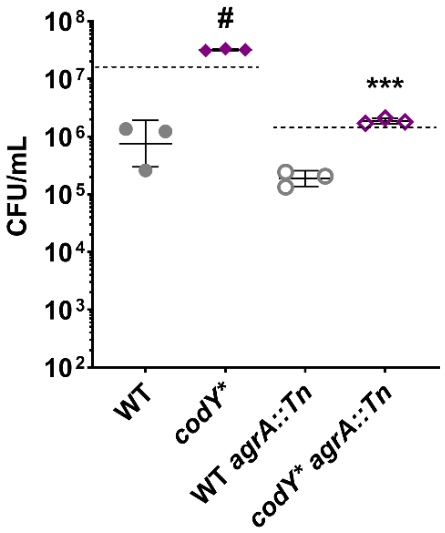

**Supplemental Figure 5. Increased PYO tolerance conferred by the *codY** mutation does not require *agr* activity**. Viable cells counts of WT, the *codY** mutant, WT *agrA*::Tn, and a *codY** *agrA*::Tn double mutant after 20-hour treatment with 200 μM PYO. Dashed lines indicate the approximate CFU at the time of PYO addition for the indicated strains. Data shown are the geometric mean ± geometric standard deviation of three biological replicates. The dashed lines indicate the mean initial cell density (CFU/mL) for the strains they overlap. Significance is shown for a comparison to the respective WT background and was determined by a one-way ANOVA using Šídák's correction for multiple comparisons. ($^{***}P < 0.001$, $^{\#}P < 0.0001$).

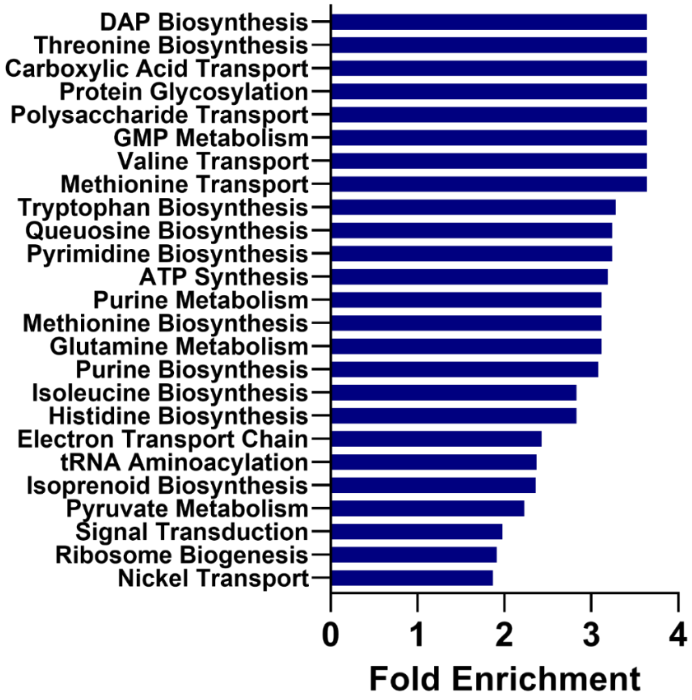

**Supplemental Figure 6. Enriched pathways among the downregulated genes in the WT response to PYO after 120 minutes overlap significantly with that of the *codY** regulon**. All enriched pathways among downregulated genes in the WT response to PYO compared to WT in DMSO after 120 minutes.

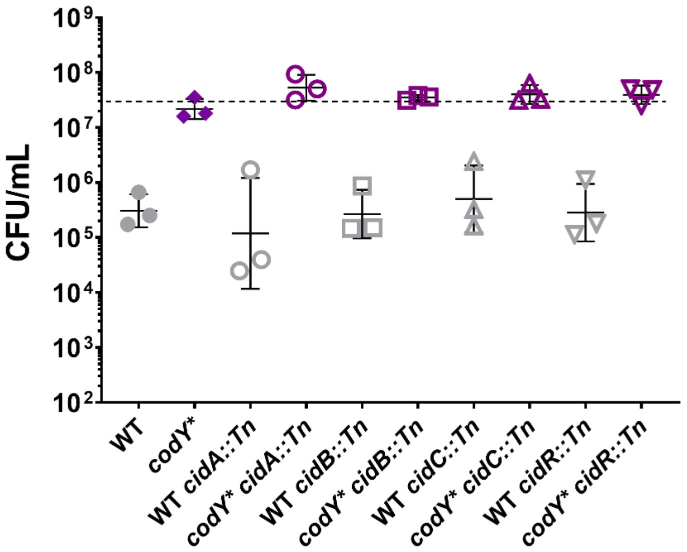

**Supplemental Figure 7. Mutations in the *cidABCR* operon do not impact PYO survival in either the WT or *codY** backgrounds**. Viable cell counts of WT, the *codY** mutant, and their respective *cid* transposon mutants after 20-hour treatment with 200 µM PYO. Data shown are the geometric mean ± geometric standard deviation of three biological replicates. The dashed line indicates the mean initial cell density (CFU/mL) for all strains. Significance was determined relative to the isogenic parental (WT or *codY**) strain by a two-way ANOVA using Dunnett's correction for multiple comparisons.

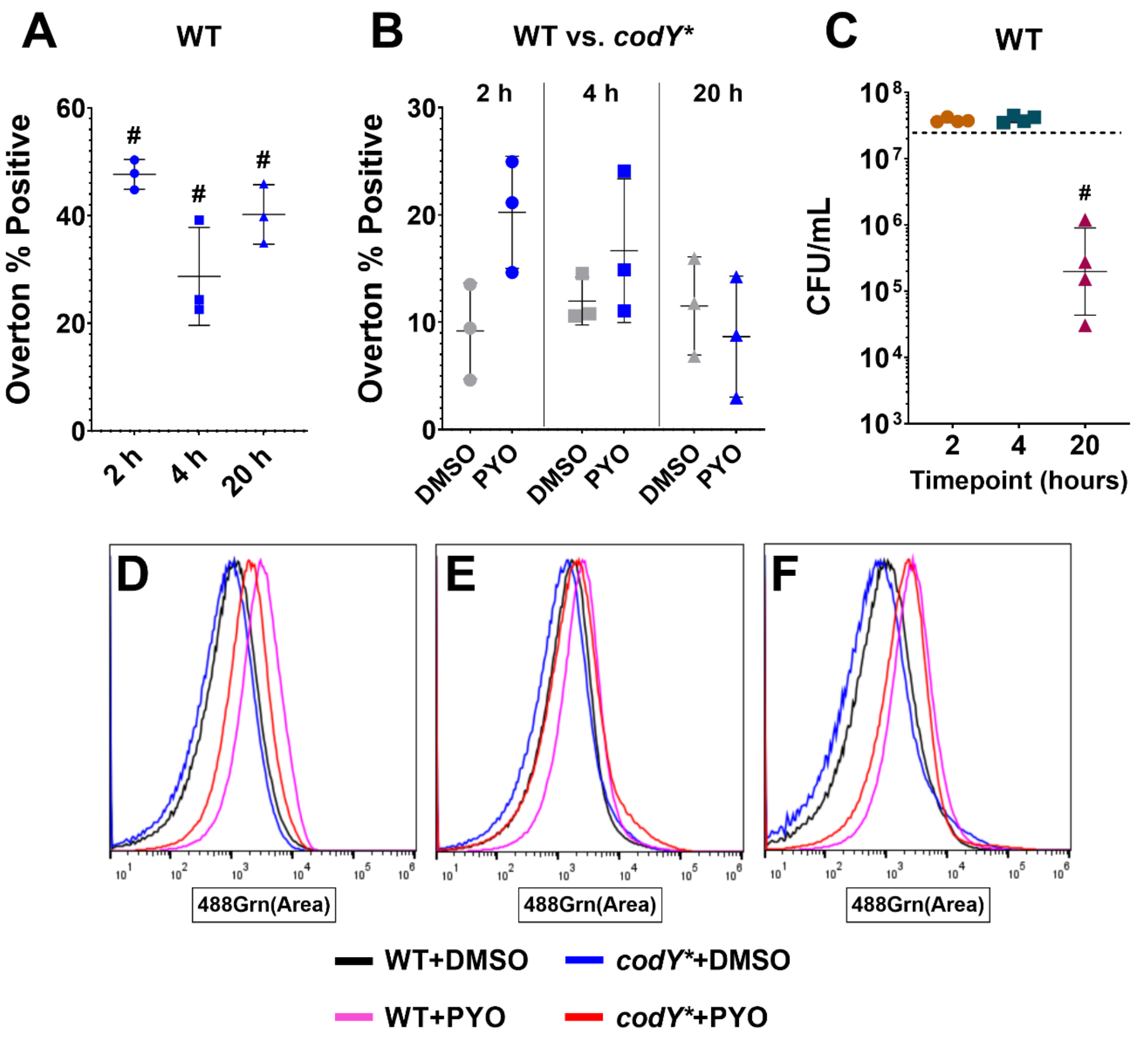

**Supplemental Figure 8. PYO induces DNA damage in *S. aureus*.** (**A**) Overton % positivity (reflecting the % of the population that has significant fluorescence compared to the control condition) after 2-, 4-, and 20-hour treatment with 200 μM PYO. (**B**) The difference in Overton % positivity between the WT and *codY** mutant in DMSO or PYO after the indicated treatment time. (**A, B**) Data shown are the mean ± standard deviation of three biological replicates. (**C**) Viable cell counts of WT after treatment with 200 μM PYO for the indicated time. Data shown are the geometric mean ± geometric standard deviation of four biological replicates. The dashed line indicates the mean initial cell density (CFU/mL). Significance is shown relative to (**A**) 0% Overton positivity, (**B**) the respective DMSO condition, or (**C**) the initial (0h) cell density as tested by a (**A,**

**B**) two-way ANOVA using (**A**) Tukey's or (**B**) Šídák's correction for multiple comparisons, or (**C**) one-way ANOVA using Dunnett's correction for multiple comparisons ($^{\#}P < 0.0001$). (**D-F**) Representative histograms from the data underlying (**B**) after (**D**) 2-, (**E**) 4-, and (**F**) 20-hour treatment.

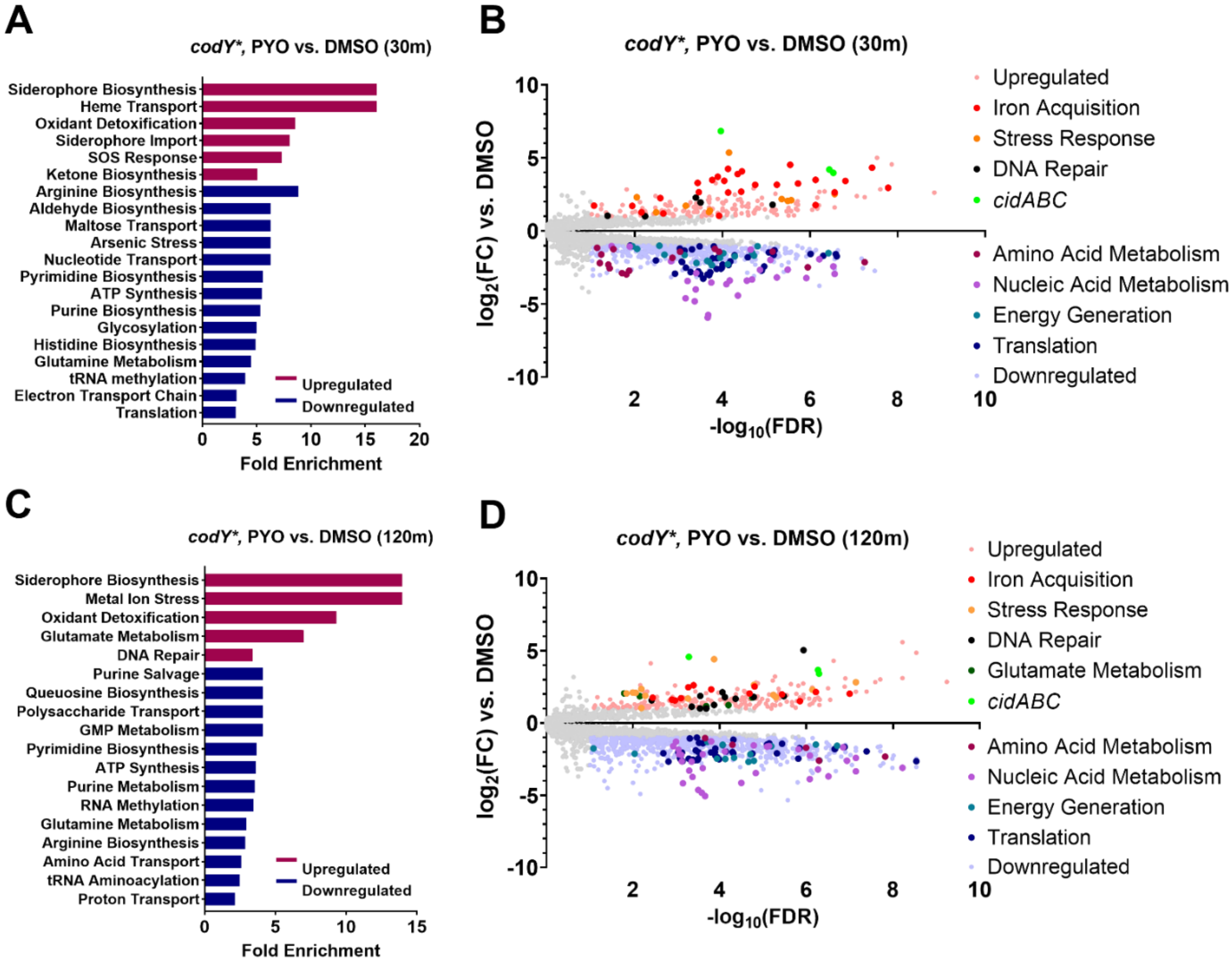

**Supplemental Figure 9. The response to PYO of the *codY** mutant is similar to that of the WT but shows greater metabolic suppression**. Transcriptional response of the *codY** mutant to PYO compared to the DMSO control after (**A**, **B**) 30 and (**C**, **D**) 120 minutes. (**A**, **C**) Enriched GO pathways from upregulated and downregulated genes. (**B**, **D**) Volcano plot of $\log_2$(fold change gene expression) and -log10(false discovery rate). Highlighted genes comprise the pathways indicated in the figure legend. A list of genes in each pathway can be found in **Supplementary Data File 03**.

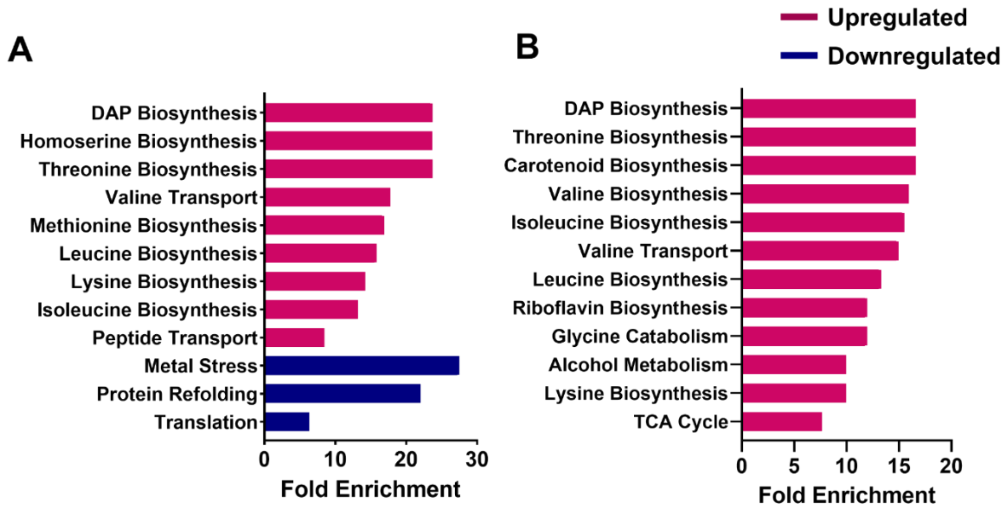

**Supplemental Figure 10. Pathways enriched within genes differentially expressed in the *codY** mutant compared to the WT in response to PYO are predominantly involved in amino acid metabolism**. Enriched GO pathways from upregulated and downregulated genes in the *codY** response to PYO compared to the WT response to PYO after 30 (**A**) and 120 (**B**) minutes of treatment with PYO.

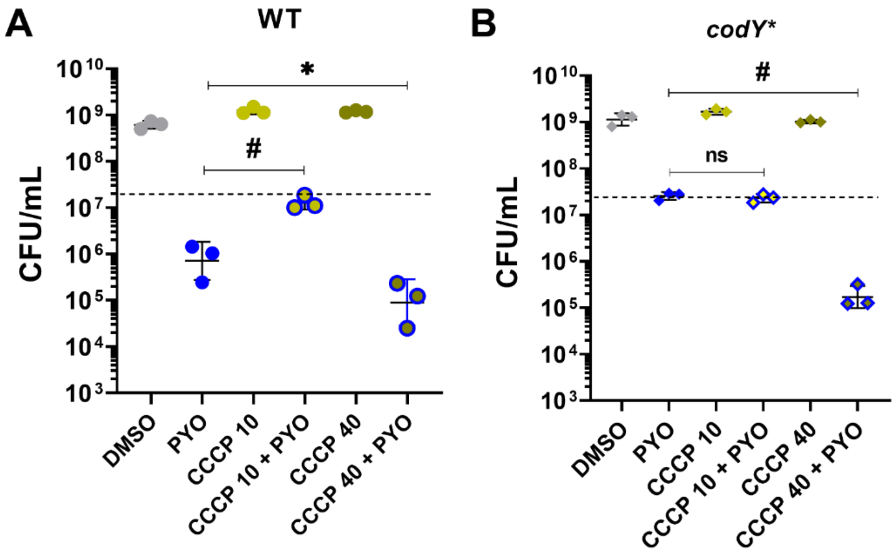

**Supplemental Figure 11. Excess CCCP sensitizes both WT and the *codY** mutant to PYO**. Effect of CCCP addition (10 μM or 40μM) on survival after 20-hour treatment with either DMSO or 200 μM PYO for the (**A**) WT and (**B**) *codY** mutant. Shown are the viable cell counts. (**A, B**) Data shown are the geometric mean ± geometric standard deviation of three biological replicates. Dashed lines indicate the mean initial cell density (CFU/mL) for all strains. Significance is shown for comparison to the indicated conditions and was determined by a one-way ANOVA using Tukey's correction for multiple comparisons. ($^{*}P < 0.05$, $^{\#}P < 0.0001$).

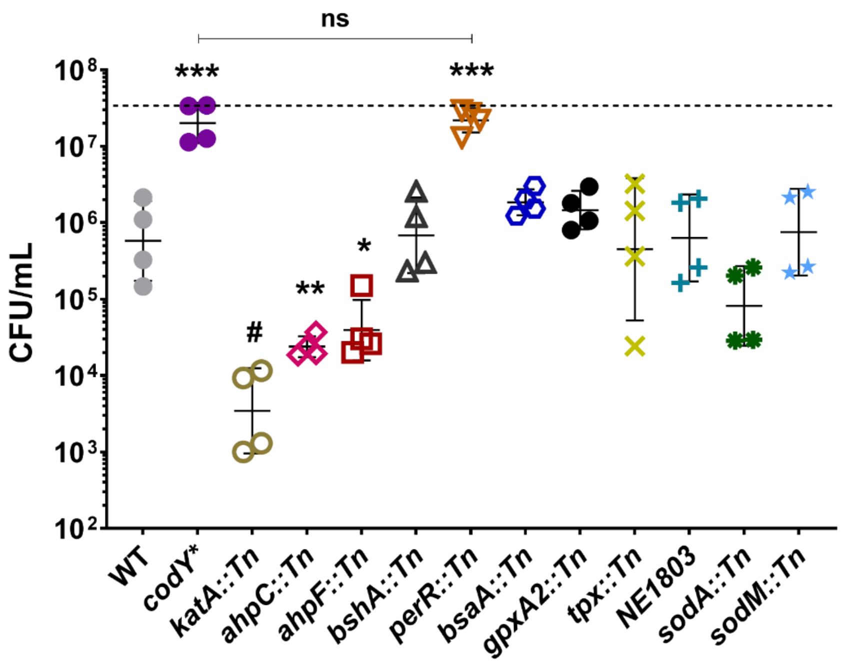

**Supplemental Figure 12. Mutants deficient in the response to hydrogen peroxide stress are more sensitive to PYO while a mutant with a constitutive stress response phenocopies *codY****. Viable cell counts of WT, the *codY** mutant, or the indicated transposon mutant strains after 20-hour treatment with 200 µM PYO. The dashed line indicates the mean initial cell density (CFU/mL) for all strains at the time of PYO addition. Data shown are the geometric mean ± geometric standard deviation of four biological replicates. Significance is indicated either for comparisons to the WT or between the indicated strains as determined by a one-way ANOVA using Dunnett's correction for multiple comparisons. ($^{*}P < 0.05$, $^{**}P < 0.01$, $^{***}P < 0.001$, $^{\#}P < 0.0001$).

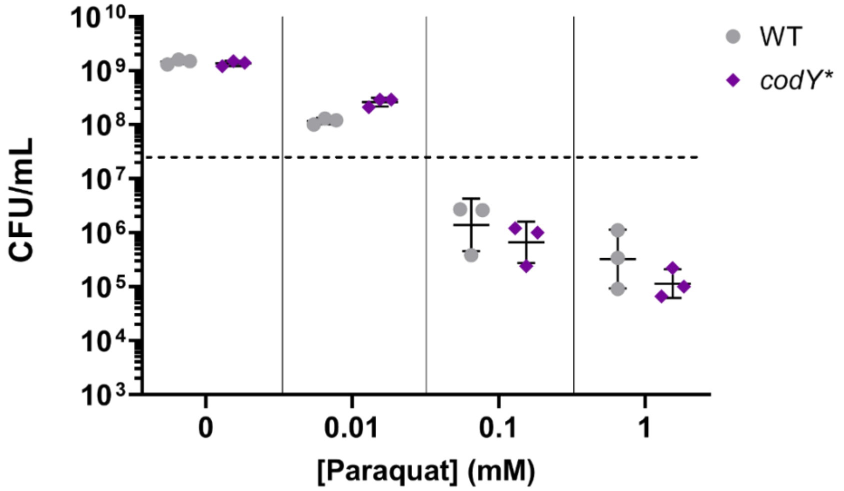

**Supplemental Figure 13. WT and the *codY** mutant are similarly susceptible to paraquat-derived superoxide**. Viable cell counts of WT and the *codY** mutant after 20-hour treatment with the indicated concentration of paraquat. Data shown are the geometric mean ± geometric standard deviation of three biological replicates. The dashed line indicates the mean initial cell density (CFU/mL) for all strains. Significance was determined for comparison to the respective WT conditions by a two-way ANOVA using Šídák's correction for multiple comparisons.

### SUPPLEMENTARY TABLES

**Supplemental Table 1. CodY-associated mutations in PYO-evolved isolates.**

| Designation | Population | Isolate | CodY Mutation |
|---|---|---|---|
| | | **Population A** | |
| 200.5-A-1 | A | 1 | Intergenic (+15/-9)<br>AG → GT |
| 200.5-A-2 | A | 2 | Intergenic (+15/-9)<br>AG → GT |
| 200.5-A-3 | A | 3 | R222C |
| 200.5-A-4 | A | 4 | S178L |
| 200.5-A-5 | A | 5 | T125I |
| 200.5-A-7 | A | 7 | R61K |
| 200.5-A-8 | A | 8 | Intergenic (+15/-9)<br>AG → GT |
| 200.5-A-9 | A | 9 | R222C |
| 200.5-A-10 | A | 10 | Intergenic (+15/-9)<br>AG → GT |
| 200.5-A-11 | A | 11 | Intergenic (+15/-9)<br>AG → GT |
| 200.5-A-12 | A | 12 | Y75C |
| | | **Population B** | |
| 200.7-B-1 | B | 1 | G118D |
| 200.7-B-3 | B | 3 | Intergenic (+16/-9)<br>G → T |
| 200.7-B-4 | B | 4 | G118D |
| 200.7-B-5 | B | 5 | G118D |
| 200.7-B-7 | B | 7 | ATG → ATA |
| 200.7-B-8 | B | 8 | G118D |
| 200.7-B-9 | B | 9 | G118D |

**Supplemental Table 2. Strains and plasmids used in this study.**

| Name | Description | Source |
|---|---|---|
| **Strains** | | |
| ***Staphylococcus aureus*** | | |
| JE2 | Plasmid-cured derivative of the USA300 strain, LAC | (1, 2) |
| RN4220 | Restriction-deficient mutant of NCTC8325-4 | (3) |
| SB523 | JE2 *codY*$^{R222C}$ | This study |
| NE1555 | *codY*::Tn | (2) |
| SB524 | JE2 Δ*qsrR* | This study |
| NE1532 | *agrA*::Tn | (2) |
| SB525 | *agrA*::Tn *codY*$^{R222C}$ | This study |
| NE1366 | *katA*::Tn | (2) |
| SB526 | *katA*::Tn *codY*$^{R222C}$ | This study |
| NE911 | *ahpC*::Tn | (2) |
| SB527 | *ahpC*::Tn *codY*$^{R222C}$ | This study |
| NE1571 | *ahpF*::Tn | (2) |
| NE1728 | *bshA*::Tn | (2) |
| NE665 | *perR*::Tn | (2) |
| NE390 | *gltB*::Tn | (2) |
| SB529 | *gltB*::Tn *codY*$^{R222C}$ | This study |
| NE445 | *umuC*::Tn | (2) |
| SB530 | *umuC*::Tn *codY*$^{R222C}$ | This study |
| NE563 | *gpxA2*::Tn | (2) |
| SB531 | *gpxA2*::Tn *codY*$^{R222C}$ | This study |
| NE1730 | *bsaA*::Tn | (2) |
| SB532 | *bsaA*::Tn *codY*$^{R222C}$ | This study |
| NE1803 | RS04260::Tn | (2) |
| NE1332 | *tpx*::Tn | (2) |
| NE1932 | *sodA*::Tn | (2) |
| NE1224 | *sodM*::Tn | (2) |
| NE122 | *adhC*::Tn | (2) |
| SB533 | *adhC*::Tn *codY*$^{R222C}$ | This study |
| NE1538 | *crtO*::Tn | (2) |
| SB534 | *crtO*::Tn *codY*$^{R222C}$ | This study |
| NE1692 | *cidA*::Tn | (2) |
| SB536 | *cidA*::Tn *codY*$^{R222C}$ | This study |
| NE935 | *cidB*::Tn | (2) |
| SB537 | *cidB*::Tn *codY*$^{R222C}$ | This study |
| NE564 | *cidC*::Tn | (2) |
| SB538 | *cidC*::Tn *codY*$^{R222C}$ | This study |
| NE1466 | *cidR*::Tn | (2) |
| SB539 | *cidR*::Tn *codY*$^{R222C}$ | This study |
| ***Escherichia coli*** | | |
| DC10B | Cloning *E. coli* DC10B, *dam*/*dcm*-deficient | (4) |
| SB192 | Cloning *E. coli* IM08B with USA300-pattern restriction modification | (5) |

| | | |
|---|---|---|
| **Plasmids** | | |
| pIMAY* | Allelic exchange vector | (6) |
| pIMAY*-codY$^{R222C}$ | Allelic exchange vector for construction of the *codY** allele | This study |
| SB209 | pKM16; fluorescent reporter | (7) |
| SB540 | pKM16 backbone expressing *katA* from its native promoter | This study |
| SB541 | pKM16 backbone expressing *ahpCF* from its native promoter | This study |
| SB542 | pKM16 backbone expressing *umuC* from its native promoter | This study |
| SB543 | pKM16 backbone expressing *pxpBCA* from its native promoter | This study |
| SB544 | pKM16 backbone expressing *adhC* from its native promoter | This study |

**Supplemental Table 3. Oligonucleotides used in this study.**

| Primer Name | Description | Primer Sequence* |
|---|---|---|
| Sa035 | Forward amplification of *codY* adding an EcoRI site with homologous sequence to pIMAY* for Gibson assembly | GGTATCGATAAGCTTGATATC**GAATTC**CGACAAAGTTGCGACGAATA |
| Sa036 | Reverse amplification of *codY* adding a NotI site with homologous sequence to pIMAY* for Gibson assembly | TGGAGCTCCACCGCGGTG**GCGGCCGC**ATCACGAATTCCGCCTAAGA |
| Sa037 | Forward pIMAY*-*codY* integration primers | AAGAAAGTTGAGCGAGAA |
| Sa038 | Reverse pIMAY*-*codY* integration primers | AATTGCATTACGCTCTTT |
| JE2/qsrR/Up.F1 | Upstream fragment for *qsrR* allelic exchange | TCACTAAAGGGAACAAAAGCTCTTTAACAACTAAATCATCTCCG |
| JE2/qsrR/Up.R1 | Upstream fragment for *qsrR* allelic exchange | AGATGATGGAAGTATGTCCGCGTACTGCTAAATAATTACATGACG |
| IMAY*/qsrR/Up.F2 | Vector linearization for *qsrR* allelic exchange | GCCTGCTTCAATTCTATCAGCGTCACAGGTATTTATTCGG |
| IMAY*/qsrR/Up.R2 | Vector linearization for *qsrR* allelic exchange | GATGATTTAGTTGTTAAAGAGCTTTTGTTCCCTTTAGTGA |
| JE2/qsrR/Down.F3 | Downstream fragment for *qsrR* allelic exchange | TGTAATTATTTAGCAGTACGCGGACATACTTCCATCATCTTC |
| JE2/qsrR/Down.R3 | Downstream fragment for *qsrR* allelic exchange | CCGAATAAATACCTGTGACGCTGATAGAATTGAAGCAGGC |
| NTML/Buster2.R | Transposon primer (reverse) | GCCAACCTGTTACTAGACCG |
| NTML/Upstream2.F | Transposon primer (forward) | AAAGCATTGAACACCATAACCG |
| JE2/agrA/NTML.R1 | Screening of *agrA* transposon insertion | TACACTGAATTACTGCCACG |
| JE2/cidA/NTML.R1 | Screening of *cidA* transposon insertion | GAAAATGAAGTGAAATTTAGAGAGC |
| JE2/cidB/NTML.F1 | Screening of *cidB* transposon insertion | TCATCCCAATTGAACTAAATGC |
| JE2/cidC/NTML.F1 | Screening of *cidC* transposon insertion | GTTTAAGAATGGTCTTTCAGCA |
| JE2/cidR/NTML.R1 | Screening of *cidR* transposon insertion | GAAATAGTTAGGATGATGTTAGTGG |
| JE2/katA/NTML.F1 | Screening of *katA* transposon insertion | GCAGCTTGTTCAACATCC |
| JE2/ahpC/NTML.R1 | Screening of *ahpC* transposon insertion | CATTACCTTCATCCATCTCG |
| JE2/ahpF/NTML.F1 | Screening of *ahpF* transposon insertion | GCACGTATACCTGTCATTGC |
| JE2/perR/NTML.F1 | Screening of *perR* transposon insertion | ATCATTGCGACAAGCAGG |
| JE2/gltB/NTML.F1 | Screening of *gltB* transposon insertion | CTTGAAATGTTGCGACGC |

| | | |
|---|---|---|
| JE2/gpxA2/NTML.F1 | Screening of *gpxA2* transposon insertion | TCATTAGAACCTGGTTGTCG |
| JE2/bshA/NTML.F1 | Screening of *bshA* transposon insertion | GATCATTCACTCCAAGGTGC |
| JE2/tpx/NTML.F1 | Screening of *tpx* transposon insertion | ATAACATTCAAAGGTGGACC |
| JE2/umuC/NTML.F1 | Screening of *umuC* transposon insertion | GTTGTTGCAGATACTAAGCG |
| JE2/RS04260/NTML.R1 | Screening of *RS04260* transposon insertion | TTTATGGATTGACCCAATCG |
| JE2/crtO/NTML.R1 | Screening of *crtO* transposon insertion | TCTCTAACATGAGAGTATTGGC |
| JE2/adhC/NTML.R1 | Screening of *adhC* transposon insertion | ATACCTCCTAAACCAACAACCGC |
| pKM16/Dn/Linear.F1 | Linearization of pKM16 excluding *sarA_dsRed* | CGGTTATCCACAGAATCAGG |
| pKM16/Up/Linear.R1 | Linearization of pKM16 excluding *sarA_dsRed* | AAAGATCCTAACGAAAAGCG |
| JE2/umuC/Up.F1 | *umuC* upstream primer for pKM_umuC | CGCTTTTCGTTAGGATCTTTCGATTGGCAACATCCAAACC |
| JE2/umuC/Dn.R1 | *umuC* downstream primer for pKM_umuC | CCTGATTCTGTGGATAACCGAAACCCTACTGACCGAGAAC |
| JE2/katA/Up.F1 | *katA* upstream primer for pKM_umuC | CGCTTTTCGTTAGGATCTTTTAAAATGTTGCCAACTCTCC |
| JE2/katA/Dn.R1 | *katA* downstream primer for pKM_umuC | CCTGATTCTGTGGATAACCGCATAAACTGCTCAACTACGC |
| JE2/ahpCF/Up.F1 | *ahpCF* upstream primer for pKM_umuC | CGCTTTTCGTTAGGATCTTTATCTTCTCATCGTCGATACC |
| JE2/ahpCF/Dn.R1 | *ahpCF* downstream primer for pKM_umuC | CCTGATTCTGTGGATAACCGAAGCATTATCGCACATCTCG |
| JE2/pxpA/Up.F1 | *pxpA* upstream primer for pKM_umuC | CCTGATTCTGTGGATAACCGTTCTCCCCATTTTTTTAGCC |
| JE2/pxpA/Dn.R1 | *pxpA* downstream primer for pKM_umuC | GAAAAAATCGATGCGAGTTGATTTGAATTG |
| JE2/pxpBC/Up.F1 | *pxpBC* upstream primer for pKM_umuC | CAACTCGCATCGATTTTTTCAATATTGATTTTACAAATCC |
| JE2/pxpBC/Dn.R1 | *pxpBC* downstream primer for pKM_umuC | CGCTTTTCGTTAGGATCTTTGAATCACCAATGGCTAAAGC |
| JE2/adhC/Up.F1 | *adhC* upstream primer for pKM_umuC | CGCTTTTCGTTAGGATCTTTAGTTTCATAATCCCACTCCC |
| JE2/adhC/Dn.R1 | *adhC* downstream primer for pKM_umuC | CCTGATTCTGTGGATAACCGGAGCATCCTTCACTTTTGCG |

***Bold indicates the restriction cut site.**

**Underlined sequences indicate regions of homology for Gibson assembly.

### SUPPLEMENTARY DATA FILES

**Supplementary Data File 01: A list of all mutations observed in each sequenced evolved isolate.**

**Supplementary Data File 02: The full edgeR analysis, differentially expressed genes, and enriched PANTHER pathways for each RNA-seq comparison.**

**Supplementary Data File 03: A list of genes comprising each pathway shown in the legend of the volcano plots.**

**Supplementary Data File 04: CodY sequences of *S. aureus* isolates shown by position.**

**Supplementary Data File 05: Scripts used for RNA-seq alignment and edgeR analysis.**

**Supplementary Data File 06: Raw processed data files from kallisto in tsv format.**
